# Long-term changes in the timing of autumn migration in Alaska’s boreal songbirds

**DOI:** 10.1101/2023.09.05.556417

**Authors:** April Harding Scurr, Julie Hagelin, Grey Pendleton, Kristin DuBour, Tricia Blake, Claire Stuyck, Eva Allaby

**Affiliations:** Alaska Songbird Institute, Fairbanks, Alaska, United States of America; Threatened, Endangered, and Diversity Program, Alaska Department of Fish and Game, Fairbanks, Alaska, United States of America; Threatened, Endangered, and Diversity Program, Alaska Department of Fish and Game, Juneau, Alaska, United States of America; National Wildlife Refuge System Inventory & Monitoring, U.S. Fish and Wildlife Service, Anchorage, Alaska, United States of America

## Abstract

Alaska’s boreal birds face a rapidly changing environment, but we know little about shifts in migratory timing, particularly in autumn. We used quantile regression to quantify long-term changes in autumn capture date in 21 boreal passerines using 22+year datasets from two banding stations in central Alaska. We also quantified differences between sites and explored whether select climate indices during three periods of the annual cycle (breeding, post-fledge, and migration) could predict long-term changes in median capture. Long-term changes in autumn migration were detected in 86% of taxa, 76% of which exhibited advances in capture date (∼2-3 days/decade), particularly long-distance migrants at one field site. However, site-specific differences unexpectedly highlight the need for caution before extrapolating long-term timing patterns over broad spatial extents. Warmer conditions during the breeding period (using the AO climate index) were associated with advances in autumn capture date in the greatest number of species (9). Collectively, we hypothesize that Alaska’s immense size and spatially-variable climate regions impact reproductive timing, often resulting in long-term advances (with warming) and occasionally delays (with cooling). Carry-over effects of reproductive timing may therefore influence the autumn passage of different breeding populations, causing site-specific patterns, such as a species showing long-term advances at one location, but delays at another. Finally, as part of the broader effort to anticipate and reduce declines in boreal migratory birds, our study underscores the conservation value of banding station data in quantifying avian responses to and investigating drivers associated with varied climate indices.

## INTRODUCTION

Of the estimated 2.5 billion native migratory birds lost in North America since 1970, the boreal forest biome has exhibited one of the most dramatic declines (Rosenberg et al. 2019). Half of all boreal bird species are decreasing, with a reduction of 500 million birds in the last 50 years (Rosenberg et al. 2019). The boreal forest provides critical breeding and migratory habitat for hundreds of bird species, one-third of which rely on it to sustain ≥50% of their breeding populations (Wells 2011, Wells et al. 2014). Alaska, located at the “global breeding endpoint” of nearly every major migratory flyway in North America (Kessel and Gibson 1978), is comprised of 60% boreal forest habitat (Malone et al. 2009). Consequently, the conservation of Alaska’s boreal forest biome is critical to landbird conservation and management in North America, despite receiving less attention than Canada (Matsuoka et al. 2019).

Climate change within the boreal biome is accelerated relative to the rest of the earth (Loarie et al. 2009), with temperatures in Alaska warming at 1.5 times the rate of the continental U.S. (Ballinger et al. 2023) and twice the global average (Markon et al. 2018). Climate warming influences the timing of snowmelt and insect emergence, alters vegetation and precipitation, can change large-scale climate patterns, and increases the risk of extreme weather events (Stone et al. 2002, Tulip and Schekkerman 2008, Euskirchen et al. 2009, Markon et al. 2018). These changes are predicted to disproportionately affect boreal forest and arctic avian species (Bateman et al. 2020) and, in Alaska, may impact ranges, breeding distributions, and timing of annual life cycles (Handel et al. 2021, Wu et al. 2022). Understanding and addressing the impacts of widespread stressors such as climate is necessary for the effective management of Alaska’s boreal birds (Handel et al. 2021).

Currently, it is unclear whether migratory birds in boreal environments can alter their life cycle events sufficiently to keep pace with climate change (e.g., Visser and Both 2005, Carey 2009, Handel et al. 2017). Climate conditions during spring and summer, for example, can influence when nesting ends and migration begins (Jenni and Kéry 2003, Gordo 2007). However, in arctic and sub-arctic Alaska, conditions are only snow-free for a few months each year, creating a relatively narrow window of time for migratory birds to synchronize reproduction with an irruptive pulse of food (Benson and Winker 2001, Carey 2009, Handel et al. 2017). Therefore, the climate during the breeding period could have a magnified impact on reproductive timing and subsequent autumn migration of Alaska birds, compared to lower latitudes (Carey 2009, Benson and Winker 2015). Timing adjustments within the narrow reproductive window also carry the risk of overlapping with energetically demanding events (e.g., breeding, molt; Benson and Winker 2015, Tomotani et al. 2016), although some species show surprising resilience in timing to extreme conditions (e.g., spring weather Mizel et al. 2017).

Quantifying shifts in migratory timing and relating them to climate indices are crucial first steps toward identifying possible drivers of decline and associated conservation or management actions aimed at reducing bird losses (e.g., Knudsen et al. 2011; Mayor et al. 2017; Horton 2020, 2023). Recent investigations of some Alaska species indicate that the date of spring migration is shifting earlier (advancing; Ward et al. 2016, Mayor et al. 2017, Zaifman et al. 2017), consistent with continental patterns (Carey 2009, Mayor et. 2017, Barton and Sandercock 2018, Horton et al. 2020). Autumn migration, however, has received relatively less attention, especially in temperate and arctic ecosystems (Gallinat et al. 2015, Seavy et al. 2018), compared to eastern North America (e.g, Smith and Patton 2011, Zelt et al. 2017, Stegman et al. 2017). Alaska, in particular, remains notably understudied (Seavy et al. 2018) with only one investigation focusing on long-term changes in autumn migration timing (Zaifman et al. 2017).

We lack a general understanding of autumn migration patterns relative to climate change in Alaska, in part due to a lack of data and breeding records at high latitudes (Ward et al. 2016). In this study, we examined two long-term (20+ years) banding datasets of migrating songbirds from boreal forest habitat in central Alaska. Our primary goal was to quantify changes (advances or delays) in autumn migratory phenology for 21 passerine species, 17 of which are either a Species of Greatest Conservation Need in the State of Alaska (ADFG 2015), or listed on other national or international conservation watchlists (Table 1). Though patterns of migratory timing in autumn tend to be more variable than in spring (e.g., Horton et al. 2020), we predicted that long-distance migratory species would show more evidence for autumn advancement, consistent with other investigations (e.g., Jenni and Kéry 2003, Van Buskirk et al. 2009, Gallinat et al. 2015, Zaifman et al. 2017). We also predicted that any patterns of advancement or delays would be similar at each banding site, due to their close proximity (277 km apart) and similar habitats. Like Barton and Sandercock (2018), we used the capture date as a proxy for migration timing and calculated long-term trends for each quantile of the autumn passage period. Finally, we also explored how advances or delays in the median migration date of Alaska birds correlated with multiple, broad-scale climate indices (Barton and Sandercock, 2018).

**Table 1:** Avian taxa included in our analysis of long-term trends in timing of autumn migration. The specific state, federal or international conservation “watchlists” associated with particular species. Species names followed by a solid circle are long-distance migrants, and those with open triangles are short-distance migrants (Billerman et al. 2022).

| Common Name | Scientific Name | Species Code |
| --- | --- | --- |
| Alder Flycatcher •,<sup>1</sup> | *Empidonax alnorum* | ALFL |
| Hammond’s Flycatcher• | *Empidonax hammondii* | HAFL |
| Gray-cheeked Thrush •,2 | *Catharus minimus* | GCTH |
| Swainson's Thrush •,1 | *Catharus ustulatus* | SWTH |
| Orange-crowned Warbler •,1,2,3 | *Oreothlypis celata* | OCWA |
| Yellow Warbler •,1,3 | *Setophaga petechia* | YEWA |
| Yellow-rumped Warbler• | *Setophaga coronata* | YRWA |
| Blackpoll Warbler •,1,3 | *Setophaga striata* | BLPW |
| Northern Waterthrush• | *Parkesia noveboracensis* | NOWA |
| Wilson's Warbler •,1,2 | *Cardellina pusilla* | WIWA |
| Savannah Sparrow •,1 | *Passerculus sandwichensis* | SAVS |
| Lincoln's Sparrow •,1 | *Melospiza lincolnii* | LISP |
| Ruby-crowned Kinglet<sup>Δ,1</sup> | *Regulus calendula* | RCKI |
| Hermit Thrush <sup>Δ,1</sup> | *Catharus guttatus* | HETH |
| American Robin<sup>Δ</sup> | *Turdus migratorius* | AMRO |
| Varied Thrush <sup>Δ,1,2</sup> | *Ixoreus naevius* | VATH |
| American Tree Sparrow <sup>Δ,1,2</sup> | *Spizella arborea* | ATSP |
| Fox Sparrow <sup>Δ,1,2,3</sup> | *Passerella iliaca* | FOSP |
| White-crowned Sparrow <sup>Δ,1,2</sup> | *Zonotrichia leucophrys* | WCSP |
| Dark-eyed Junco <sup>Δ,1,2</sup> | *Junco hyemalis* | DEJU |
| Rusty Blackbird <sup>Δ,4,5,6</sup> | *Euphagus carolinus* | RUBL |

<sup>1</sup>Alaska Department of Fish and Game Species of Greatest Conservation Need (ADFG 2015)
<sup>2</sup>Partners in Flight Continental Stewardship Species – a species with ≥10% of its global
population occurring in North America and ≥25% of its North American population occurring in
Alaska (Panjabi et al. 2020, Handel et al. 2021)
<sup>3</sup>Audubon Alaska Watchlist 2017 (Warnock. 2017).
<sup>4</sup>Bureau of Land Management Special Status Species (BLM 2019)
<sup>5</sup>Department of Defense Mission Sensitive Species (DOD PIF 2021)
<sup>6</sup>IUCN Vulnerable Species (IUCN 2021)

## METHODS

### Study sites

We used more than 20 years of autumn migration data from two passerine banding stations (Figure 1), located 277 km apart along the same major migratory flyway in eastern Interior Alaska, the Tanana River Valley (Cooper and Ritchie 1995, Sivakumar et al. 2021). The valley is a well-documented bird migration corridor (Kessel 1984, McIntyre and Ambrose 1999, Benson and Winker 2001) and, for this reason, the Upper River Tanana Valley is recognized as a Global Important Bird Area (Smith et al. 2014, BirdLife International 2023). Data collection occurred at Creamer’s Field Migration Station (CFMS; 1992-2018), within Creamer’s Field Migratory Waterfowl Refuge, Fairbanks Alaska, (64.86754° N, 147.7479° W, elevation: 136 m), and at a former U.S. Army fuel pumping station (PUMP; 1993-2015) located seven miles west of Tok, Alaska near Tetlin National Wildlife Refuge and operated by the U.S. Fish and Wildlife Service (63.36177° N, 143.212° W; elevation 494 m).

**Figure 1:**
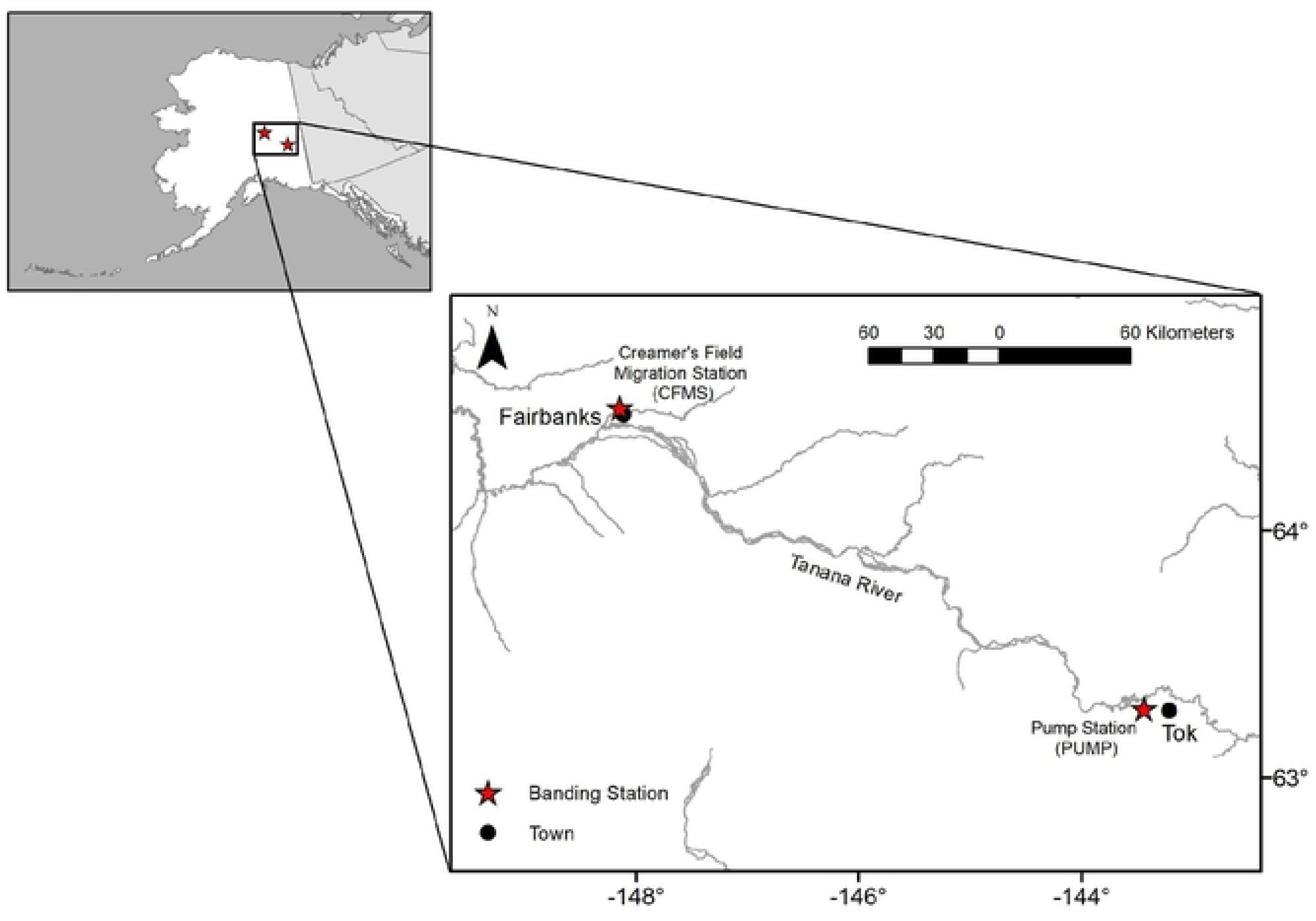
The location of the Creamer’s Field Migration Station and the PUMP station, both of which are located in boreal forest habitat along the Tanana River Valley migratory flyway in central Alaska, U.S.A.

Both banding stations have undergone a similar pattern of successional change and share a variety of habitats and species common to Alaska’s interior boreal forest. Common trees include quaking aspen (*Populus tremuloides*), Alaska birch (*Betula neoalaskana*), balsam poplar (*Populus balsamifera*), white spruce (*Picea glauca*), and common understory species include willows (*Salix* sp.), Siberian alder (*Alnus viridis*), sedges (*Carex* spp.), and bluejoint grass (*Calamagrostis canadensis*). CFMS covers ∼20 ha and was a cleared, working dairy farm until 1966. Its lands were preserved thereafter and formally converted into a State Waterfowl Refuge in 1979 (Tauzer 2013). The northern portion of CFMS is dominated by a wet graminoid herbaceous wetland habitat (Viereck et al. 1992) bordered by Alaska birch. The southern portion is late-successional, closed mixed forest (Viereck et al. 1992) containing white spruce, balsam poplar, quaking aspen, and bordered by an agricultural field. The PUMP station covers 3 ha and was also cleared of vegetation in the 1960s during the construction of the Haines-Fairbanks Pipeline. In 1996, three years after the establishment of the PUMP banding station, the site was categorized as open/closed broadleaf forest and then closed mixed forest in 2014 (Viereck et al. 1992, USFWS 2015), consisting of deciduous and coniferous species described above. The northern edge of the PUMP station was never cleared and is dominated by mature white spruce.

### Study species

We examined long-term trends in migration timing of 21 passerine species (Table 1), based on the following criteria: (1) species were on one or more agency watchlists (Table 1), (2) individuals were captured annually at both banding sites (see Supporting Information Table S1), and (3) a sample of 20 or more individuals were captured annually at one of the study sites.

Rusty Blackbird did not meet the latter two criteria but were included in the study because of high conservation concern for this species (see Table 1). Table 1 also classifies 12 of the 21 species as “long-distance” migrants (spend non-breeding months in South America), and 9 species as “short-distance” migrants, species that spend their non-breeding months in North or Central America, (Billerman et al. 2022).

### Bird capture and long-term data collection

Bird migration data were collected at CFMS over 27 years (1992–2018) and at PUMP over 22 years (1993-2015). Nets were operated daily from August 1 through the third week of September at both sites, weather permitting. Each location employed well-established, standardized mist-netting protocols, including 20-50 mist nets (12 m x 2.6 m, 30 mm mesh) operated for at least six hours per day (Ralph 1993, Benson et al. 2012, Benson et al. 2015, USFWS 2015). CFMS collected an additional week of autumn migratory data during the last week of September when nets were operated at least every other day. In addition, only CFMS operated nets during spring migration (last week of April through the third week of May), and in some years (1992-2009), through the end of July for a breeding research project. CFMS’s spring and summer banding data helped us identify the breeding period of each species for analyses of how climate may correlate with autumn migration (see: Influence of climate on capture timing).

Upon capture, each bird was identified to species and received a federal aluminum band. We categorized each captured individual as either Hatch Year (HY; hatched that calendar year), After Hatch Year (AHY; hatched during a previous calendar year), and determined its sex (whenever possible), based on standard methods described in Pyle (1997, 2022), such as plumage, molt, degree of skull pneumatization, and breeding status (e.g., incubation patches in females, enlarged cloacal protuberances in males). We did not include HY individuals with juvenile body plumage ≥ ⅔ retained, which indicated locally fledged young that were not actively migrating (Pyle 1997, 2022). Each spring, the first capture or observation at CFMS was recorded as the first arrival date for a species.

Of the 27 years and 22 years of autumn banding records from CFMS and PUMP, respectively, we included birds captured starting August 1st. We used August 1 because previous work indicated that local breeding was usually complete at both banding sites by this date (Benson et al. 2012). Bird data on or after this date also rarely showed evidence of breeding (e.g., brood patches, cloacal protuberance) or fledging (≥ ⅔ juvenile plumage and no preformative molt). Our dataset included all newly banded birds and the first capture of returning birds banded in a previous year. We excluded birds with any evidence of non-migratory activity (e.g., breeding, recent fledging), and any birds released without bands.

The standardized netting protocol between and within seasons provided for a relatively consistent level of net effort. Therefore, we used individual captures from each banding day, rather than converting data to capture rates (i.e., the number of birds/100 net hours; [one 12 m net operated for one hour equals one net hour]). We were also interested in examining various parts of the migration distribution using quantile regression. Quantile regression was used by Barton and Sandercock (2018) and requires count data rather than capture rates.

The U.S. Fish and Wildlife project, ‘PUMP’ operated under a federal permit administered by the U.S. Geological Survey, permit number 22404; a Department of Defense land use permit; DACA85-4-00007; and the U.S. Fish and Wildlife’s Institutional Animal Care and Use Committee number 2012010. The Alaska Songbird Institute require approval for ‘CFMS’ by a designated Science Committee and also operated under a federal permit administered by the U.S. Geological Survey, permit numbers 22759 and 23850. Both research stations followed ‘Guideline to the Use of Wild Birds in Research’ (Fair et al. 2023).

### Median capture dates, differences by age class and study site

For each species we calculated the median capture date and 95% confidence interval. For species with large numbers of captures, the CI sometimes covered <1 day. We also calculated the median capture date for combinations of site and age class. We computed differences between median capture dates, between age classes, and between sites along with confidence intervals for each of these differences.

### Long-term changes in capture timing

We estimated long-term changes in the timing of autumn migration using quantile regression (Cade and Noon 2003, Knudsen et al. 2007) with 0.10, 0.25, 0.50, 0.75, and 0.90 quantiles to estimate changes in early, middle, and late capture dates. We performed separate analyses for each species with the number of daily captures as the response variable. We also calculated the median capture date (95% CI) across years by species (both sites combined), age class, and capture site. We estimated timing differences by computing the differences (in days) between median capture dates, for age classes (controlling for site) and for sites (controlling for age class); we also calculated 95% CI’s for these differences. Age-class differences were calculated as the HY median capture date minus the AHY median capture date. Site differences were calculated as the CFMS median minus the PUMP median.

### Influence of climate on capture timing

To estimate the effects of climate on the autumn capture date for each of the 21 passerines, we first defined three biologically-sensitive time periods per each species to use in climate analyses: (1) Breeding, (2) Post-Fledging, and (3) Migration (Figure 2). Defining three avian time periods at a statewide scale (i.e., over multiple populations) is a better gauge of migratory phenology because it accounts for interannual variability (Gordo and Sanz 2006, Knudsen et al. 2007, Gordo 2007), similar to statewide climate indices that we will describe below (see: Climate Indices). Species-specific dates for each time period used in the analyses are provided in Supporting Information S2 Table.

**Figure 2:**
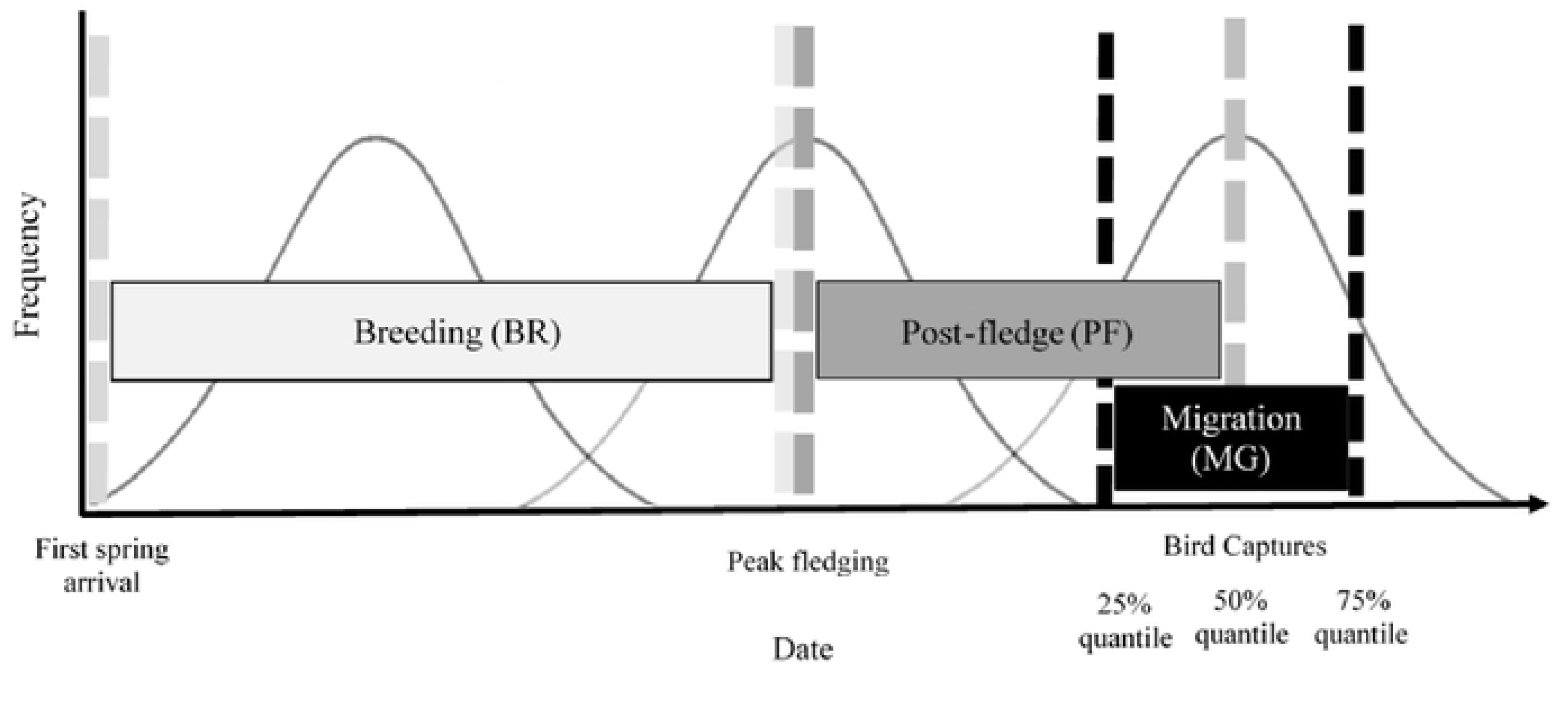
Generalized diagram representing three annual time periods (Breeding, Post-fledging, and Migration) for an example species of migratory passerine, during which it is potentially sensitive to climate impacts in Alaska. Y-axis indicates frequency (i.e., intensity) of birds statewide engaged in each period, as represented by the underlying distribution. Vertical lines indicate start and stop dates calculated for each time period (see Methods for details), to which climate indices were assigned for analysis. Species-specific dates for each period are provided in Supporting Information Table S2. Shape of frequency distributions was generalized for diagrammatic purposes and not necessarily symmetrical for all species.

We defined the Breeding Period to begin when birds first arrived in spring in central Alaska and last until peak fledging (Figure 2). To estimate the date of the first arrival in spring, we used the CFMS spring banding dataset. All available first arrival dates were averaged by species (10–14 years) over a 20-year period (1999–2018). We next assessed whether average CFMS data provided a reliable estimate for first spring arrivals statewide (e.g., whether these dates were also applicable to birds at the PUMP site). We compared CFMS average arrival dates to a range of first arrival dates for each species collected throughout central Alaska spanning 35– 41 years (1960s–1990s; Gibson 2011). CFMS estimates of first arrival fell within the species-specific date ranges of Gibson (2011) for all but 2 of the 21 species. Specifically, Varied Thrushes (Table 1) arrived 7 days later at CFMS than in Gibson (2011), and Rusty Blackbird (only 13 captures over 9 years) arrived two weeks later. In both cases, we used CFMS average dates as our spring arrival estimate for the onset of the breeding period because CFMS sample sizes were larger and better represented the banding dataset we used for analysis. Additionally, radar studies of avian migration have shown that the dates of peak spring migration to peak fall migration in Fairbanks fall very close to the average migration dates of the entire state (Sivakumar et al. 2021).

The end of the Breeding Period coincided with the onset of the Post-Fledging Period (when nestlings leave nests), which lasted until the median migratory capture date in autumn (Figure 2). Fledge dates for many passerine species in Alaska are unknown, so we used eBird’s animated weekly maps of relative abundance (2014–2019) across Alaska’s landscape to determine the week of peak fledging for each species (Fink et al. 2021). We inferred that peak statewide abundance should occur after nestlings fledge but before birds begin migrating (Morton 1991, Morton et al. 1991, Anders et al. 1998, Mitchell et al. 2010, Brown and Taylor 2015). Applying this logic to eBird relative abundance models, the Post-Fledging period began during the week when animated weekly maps of Alaska showed peak relative abundance and before the birds started moving in a predominantly southeasterly direction along the flyway representing autumn migration (Sivakumar et al. 2021). Similar to other time periods, eBird animations represented the entire state, which enabled us to calculate peak fledging dates at the statewide level. We were unable to determine an appropriate Post-Fledging period for Hermit Thrush using these methods, consequently, this species was omitted from analyses requiring the definition of a Post-Fledging period.

Finally, the Migratory Period (Figure 2) was centered around the median capture date of each species, from the 25^th^ to the 75^th^ quantile (e.g., central 50% of all autumn capture dates from both banding stations). Although the Migratory Period was the central 50% of the data distribution, the period is not necessarily symmetric about the median capture date, depending upon the distribution of the capture dates.

#### Climate indices

Climate indices account for multiple weather variables (e.g., in temperature, precipitation, wind speed) at local and regional scales (Perlwitz et al. 2017) and minimize confounding effects of intra- or interannual weather patterns (Gordo 2007), making them appropriate for understanding bird phenology (Knudsen et al. 2007). Indices are also appropriate for avian studies in Alaska, given the state’s immense size and topography, which creates a spatially variable landscape of 13 climate divisions (Bieniek et al. 2012) that potentially impact the timing of breeding and subsequent autumn migration. Since we did not know the breeding location(s) of captured birds, we selected seven climate indices that are associated with weather patterns in all climatic regions across the state (Bieniek et al. 2012, Bieniek et al. 2014; Table 2).

**Table 2.**
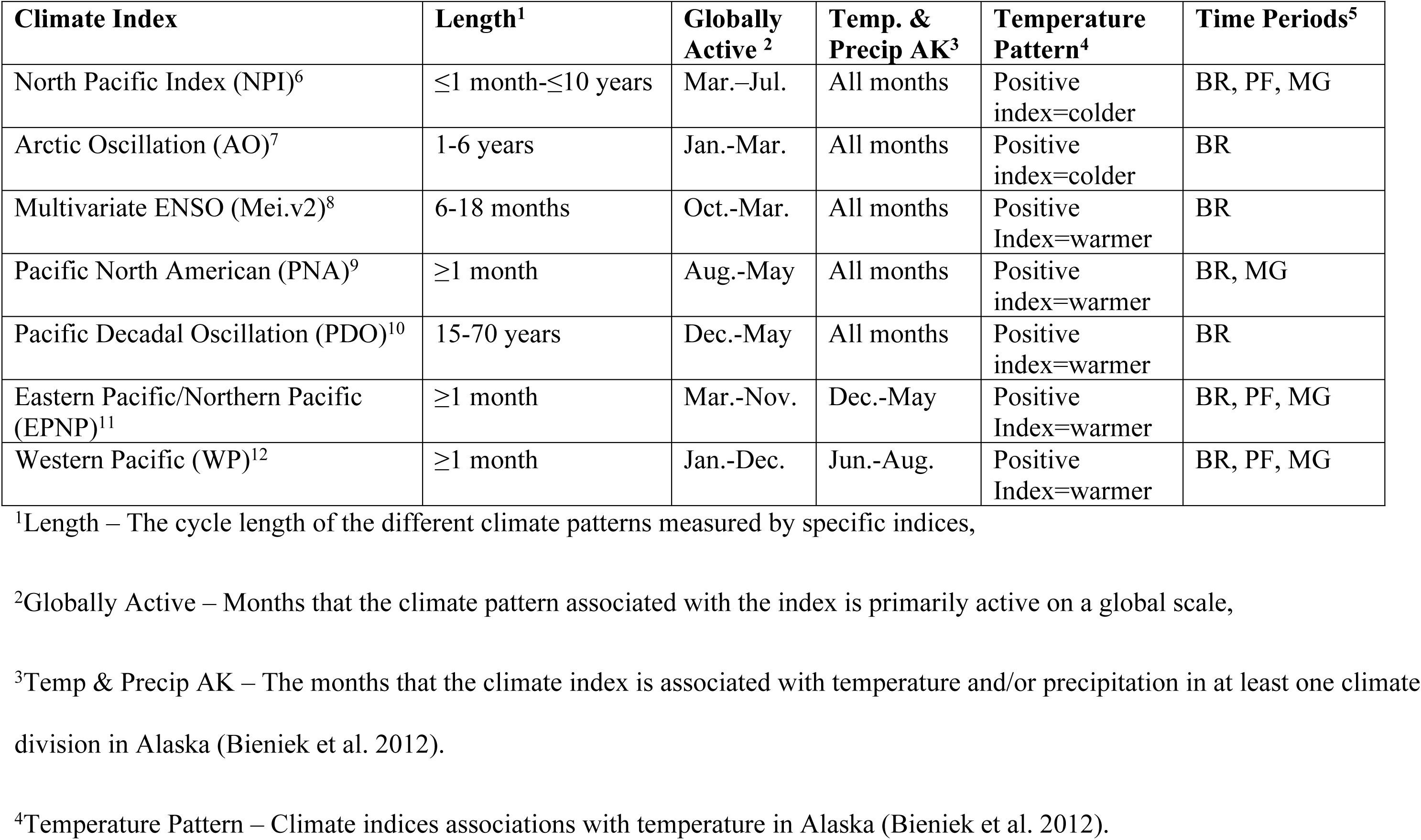
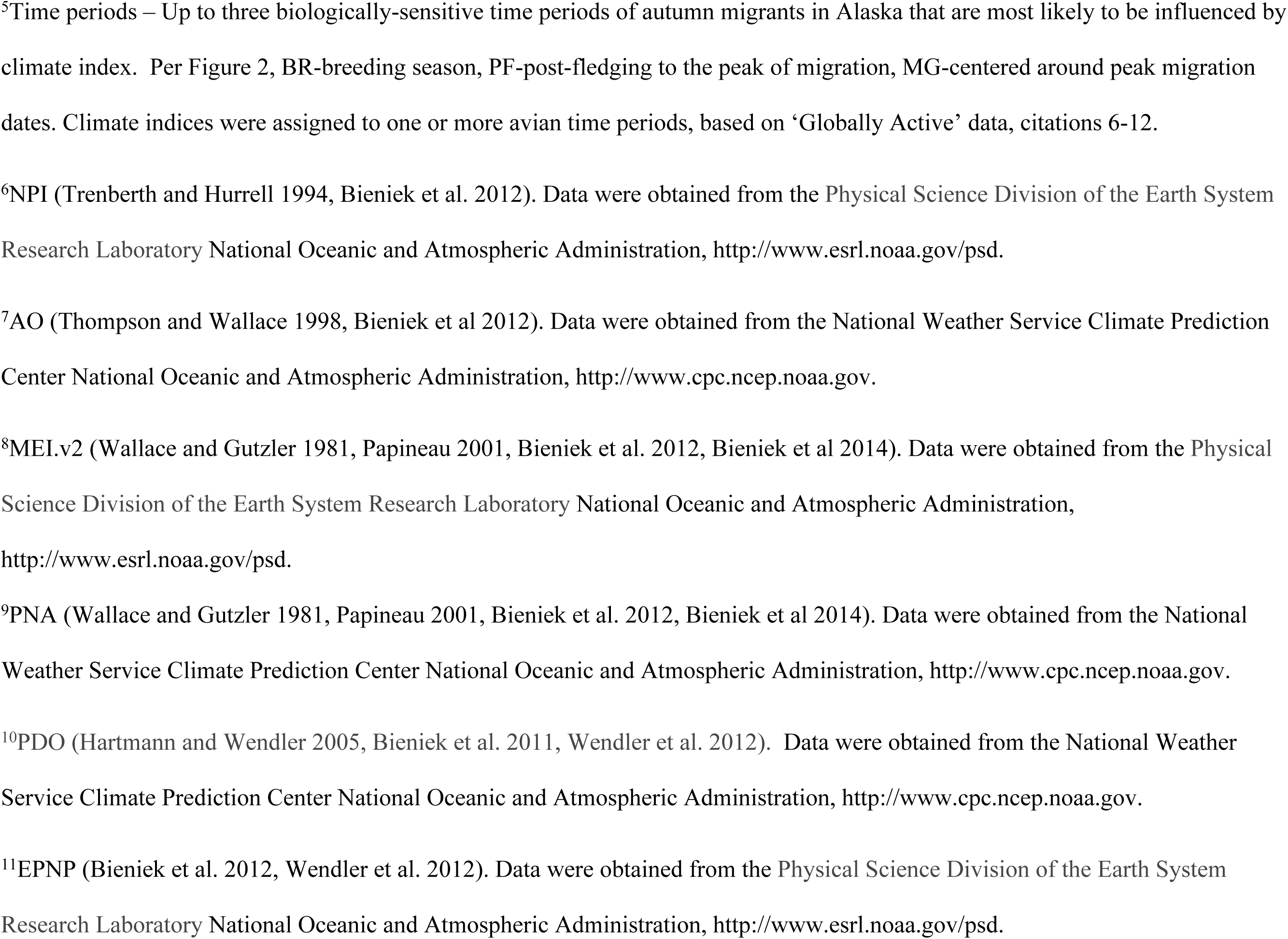
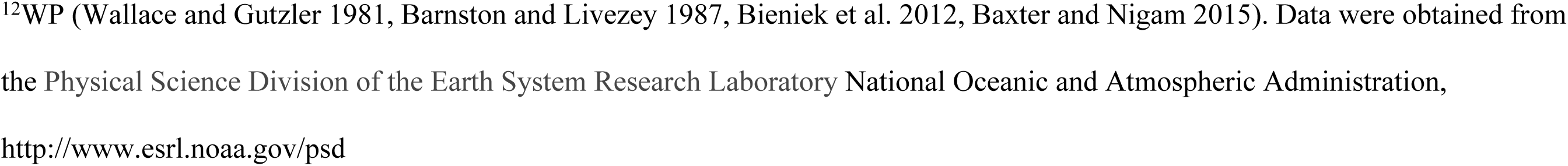
Climate indices, calculated for 3 avian life-stage periods, examined for relationships with median autumn capture dates. Although precipitation has increased 17% state-wide since 1949, patterns between specific climate indices and precipitation are still unclear (L’Heureux et al. 2004, Wendler et al. 2017). The time periods are described in Figure 2.

| **Climate Index** | **Length<sup>1</sup>** | **Globally Active<sup>2</sup>** | **Temp. & Precip AK<sup>3</sup>** | **Temperature Pattern<sup>4</sup>** | **Time Periods<sup>5</sup>** |
| --- | --- | --- | --- | --- | --- |
| North Pacific Index (NPI)<sup>6</sup> | ≤1 month-≤10 years | Mar.-Jul. | All months | Positive index=colder | BR, PF, MG |
| Arctic Oscillation (AO)<sup>7</sup> | 1-6 years | Jan.-Mar. | All months | Positive index=colder | BR |
| Multivariate ENSO (Mei.v2)<sup>8</sup> | 6-18 months | Oct.-Mar. | All months | Positive Index=warmer | BR |
| Pacific North American (PNA)<sup>9</sup> | ≥1 month | Aug.-May | All months | Positive index=warmer | BR, MG |
| Pacific Decadal Oscillation (PDO)<sup>10</sup> | 15-70 years | Dec.-May | All months | Positive index=warmer | BR |
| Eastern Pacific/Northern Pacific (EPNP)<sup>11</sup> | ≥1 month | Mar.-Nov. | Dec.-May | Positive Index=warmer | BR, PF, MG |
| Western Pacific (WP)<sup>12</sup> | ≥1 month | Jan.-Dec. | Jun.-Aug. | Positive Index=warmer | BR, PF, MG |

<sup>1</sup>Length – The cycle length of the different climate patterns measured by specific indices,
<sup>2</sup>Globally Active – Months that the climate pattern associated with the index is primarily active on a global scale,
<sup>3</sup>Temp & Precip AK – The months that the climate index is associated with temperature and/or precipitation in at least one climate division in Alaska (Bieniek et al. 2012).
<sup>4</sup>Temperature Pattern – Climate indices associations with temperature in Alaska (Bieniek et al. 2012).
<sup>5</sup>Time periods – Up to three biologically-sensitive time periods of autumn migrants in Alaska that are most likely to be influenced by climate index. Per Figure 2, BR-breeding season, PF-post-fledging to the peak of migration, MG-centered around peak migration dates. Climate indices were assigned to one or more avian time periods, based on ‘Globally Active’ data, citations 6-12.
<sup>6</sup>NPI (Trenberth and Hurrell 1994, Bieniek et al. 2012). Data were obtained from the Physical Science Division of the Earth System Research Laboratory National Oceanic and Atmospheric Administration, http://www.esrl.noaa.gov/psd.
<sup>7</sup>AO (Thompson and Wallace 1998, Bieniek et al 2012). Data were obtained from the National Weather Service Climate Prediction Center National Oceanic and Atmospheric Administration, http://www.cpc.ncep.noaa.gov.
<sup>8</sup>MEI.v2 (Wallace and Gutzler 1981, Papineau 2001, Bieniek et al. 2012, Bieniek et al 2014). Data were obtained from the Physical Science Division of the Earth System Research Laboratory National Oceanic and Atmospheric Administration, http://www.esrl.noaa.gov/psd.
<sup>9</sup>PNA (Wallace and Gutzler 1981, Papineau 2001, Bieniek et al. 2012, Bieniek et al 2014). Data were obtained from the National Weather Service Climate Prediction Center National Oceanic and Atmospheric Administration, http://www.cpc.ncep.noaa.gov.

For each of the seven climate indices (Table 2), we used published literature to determine the biologically-sensitive time periods when each index was most likely to impact, defined above. Climate patterns, as measured by the indices, operate over varying durations and annual time periods (see Table 2, footnotes 6-12). It is important to note that some indices can also have lags in Alaska, having great effects outside of the globally active months (see ‘Globally Active’ Table 2, Bieniek et al. 2012). For example, the El Nino/ La Nina Southern Oscillation climate pattern (ENSO), represented in our study by the Multivariate ENSO Index (Mei.v2), is a large and dynamic system that is a key driver of global climate that shows substantial and widespread relationships all over Alaska (Wallace and Gutzler 1981, Papineau 2001, Bieniek et al. 2012). Even though ENSO’s climate patterns are strongest from October-March globally (Table 2), in central Alaska the ENSO pattern lag behind the global pattern are primarily associated with spring temperatures that impact bird breeding, such as the timing of river-ice breakup in April or May (Bieniek et al. 2011). NPI and AO indices are scaled differently than the indices in that increases in the index indicate cooler temperatures rather than the warmers for the other indices.

#### Statistical analyses of climate impacts

We considered the seven climate indices as predictors of migration timing only during the specific time periods (Breeding, Post-Fledge, or Migration), when they were most likely to influence migration, (Table 2). Each climate index is available as monthly values. Therefore, we calculated the weighted average of monthly index values for each avian species over their specific time periods (Supporting Information Table S2). Weighted averages are based on the number of days each month contributed to the period. For example, if an avian time period included 10 days in June and 5 days in July, the calculated index value for the period would be 0.67 times the June index value + 0.33 times the July Index value.

Climate index values and median migration dates can both have linear time trends. Therefore, using climate indices to predict migration timing could result in spurious associations (Royama 1992, Van Buskirk et al. 2009). To eliminate the possibility of spurious associations, we detrended the climate indices by fitting a linear regression of each climate index on year for all analyses involving climate data. We then subtracted the predicted index value from the observed index value. The resulting residual was used as the adjusted climate index. Detrended indices were used regardless of whether the climate index by year slope had strong statistical support (i.e., 95% CI on slope included 0).

We used the adjusted climate indices (for each species and chosen time period) as predictor variables in quantile regression (Cade and Noon 2003) of the median capture date. The slope of the quantile regression is the predicted change in the median capture date for a one-unit change in the (adjusted) climate index. Climate indices, however, are not equally variable.

Consequently, we also estimated the change in the median capture date for one standard deviation of change in the (adjusted) climate index. This makes the relative effects of the climate indices on the median capture date more comparable (Barton and Sandercock 2018). Because the pattern of two climate indices (NPI and AO) are opposite of the other indices we estimate the change in median capture date for a one standard deviation *decline* in the index. For all other indices, we estimate a change of a one standard deviation *increase* in the index.

It should be noted that the climate indices are not independent and should be interpreted as a group. Also, for each index within a year, values for the adjacent time periods are calculated using data from the same months, so have an inherent correlation. Relationships among indices are most evident within each of the 3 time periods.

## RESULTS

### Bird captures and long-term data

We used 114,402 bird capture records from both study sites in our analyses: 71,907 from CFMS (27 years) and 42,495 from PUMP (22 years). Twenty species were captured at both sites and met our analysis criteria (Table 1, Supporting Information Table S1). We included one other species, Rusty Blackbird (Table 1), because of conservation concerns (see Table 1 footnotes), for a total of 21 species. Only Rusty Blackbird data from CFMS were included, as just two individuals were captured at PUMP. Although we analyzed each species by age class, we did not consider sex. Many of the 21 species we captured were monomorphic, which meant sex was undetermined for a large portion of our dataset (Supporting Information Table S1).

Median autumn capture dates varied among species from August 10 (Hammond’s Flycatcher) to September 14 (American Tree Sparrow) (Figure 3, Supporting Information Table S3). Long-distance migrants typically exhibited earlier median migration dates in autumn than short-distance migrants (Figure 3, Supporting Information Table S3). With regard to age class, median capture dates for HY birds were 6.0 days earlier (95% CI 4.4, 7.5 days) overall than median capture dates for AHY birds (17 of 21 species; Figure 4A, Supporting Information S3). The age difference pattern was consistent across sites (Figure 4A; 7.4 [4.8, 10.1] days earlier at CFMS and 5.5 [3.5, 7.4] days earlier at PUMP) Only one species (Alder Flycatcher) showed the reverse pattern with AHY median capture date occurring before HY (Supporting Information Table S3).

**Figure 3.**
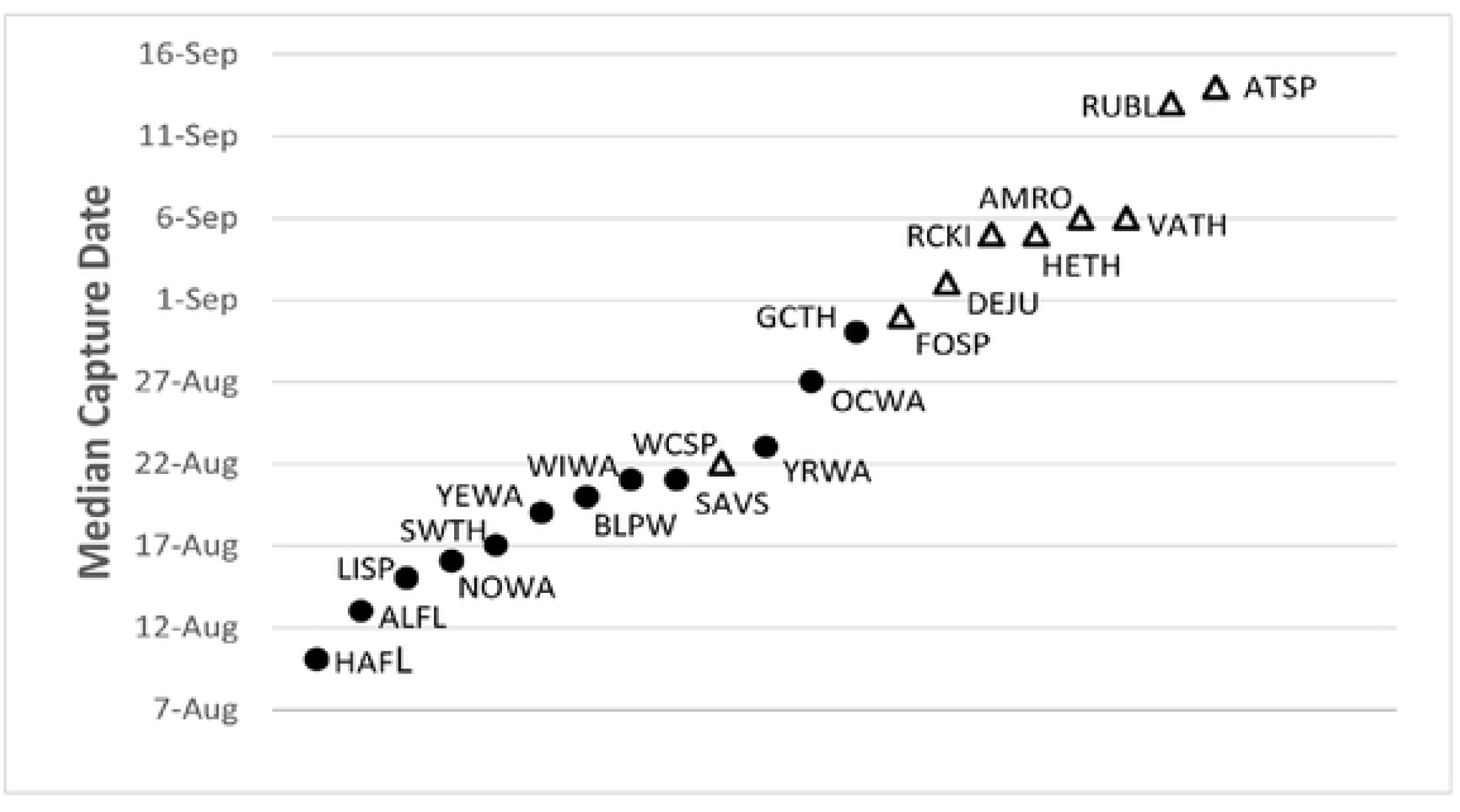
Median capture dates by species combined across years, age classes, and sites. Filled circles represent long-distance migrants and open triangles short-distance migrants.

**Figure 4.**
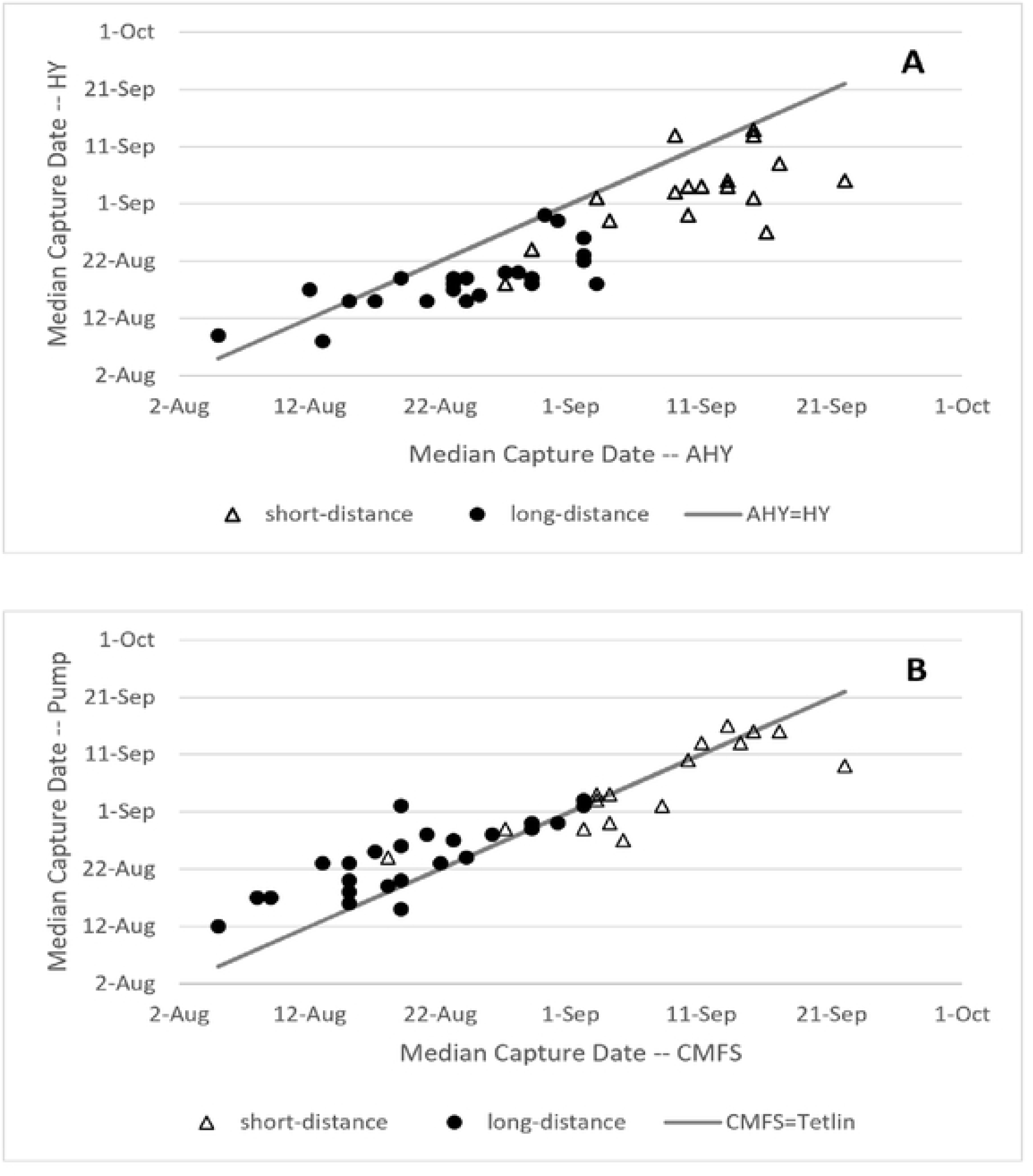
Comparison of median captures dates between ages (A) and sites (B) controlling for migratory strategy (i.e., short-distance versus long-distance migrants). Species names followed by a solid circle are long-distance migrants, and those with open triangles are short-distance migrants (Billerman et al. 2022).

Median capture dates in autumn also varied by site. Birds were generally captured earlier at CFMS than at PUMP (1.6 [-5.2, -0.0] days earlier). However, the site difference was largely driven by long-distance migrants (11 of 12 species; mean difference 3.9 [-5.6, -2.2] days), with only Swainson’s Thrush deviating from this pattern (Figure 4B, Supporting Information Table S3). We found little evidence of a consistent site difference in median capture date for short-distance migrants with two species earlier at CFMS and four species earlier at PUMP out of eight species compared (Figure 4B, Supporting Information Table S3); the mean difference in median capture date was 1.8 (-0.4, 3.4) days later at CFMS than at PUMP.

### Long-term changes in capture dates

For 18 of 21 species (86%) we found support for a long-term change in autumn capture dates (Figure 5, Supporting Information Figure S1, Table ST4). That is, the 95% CI of the estimated changes did not include 0 for one or more quantiles of autumn migratory passage (Supporting Information Figure S1, Table S4). Sixteen species (76%) showed evidence of advanced migration (earlier capture dates over time; e.g., Figures 5A, S1) in at least one quantile (of site data, or when both sites were combined). Two species exhibited long-term delays (later capture dates over time; e.g., American Tree Sparrow, Figure 5B), and three species (Northern Waterthrush, Wilson’s Warbler, and Fox Sparrow) lacked evidence of long-term changes in migration timing (Supporting Information Table S4, S5, and Supporting Information Figure S1).

**Figure 5.**
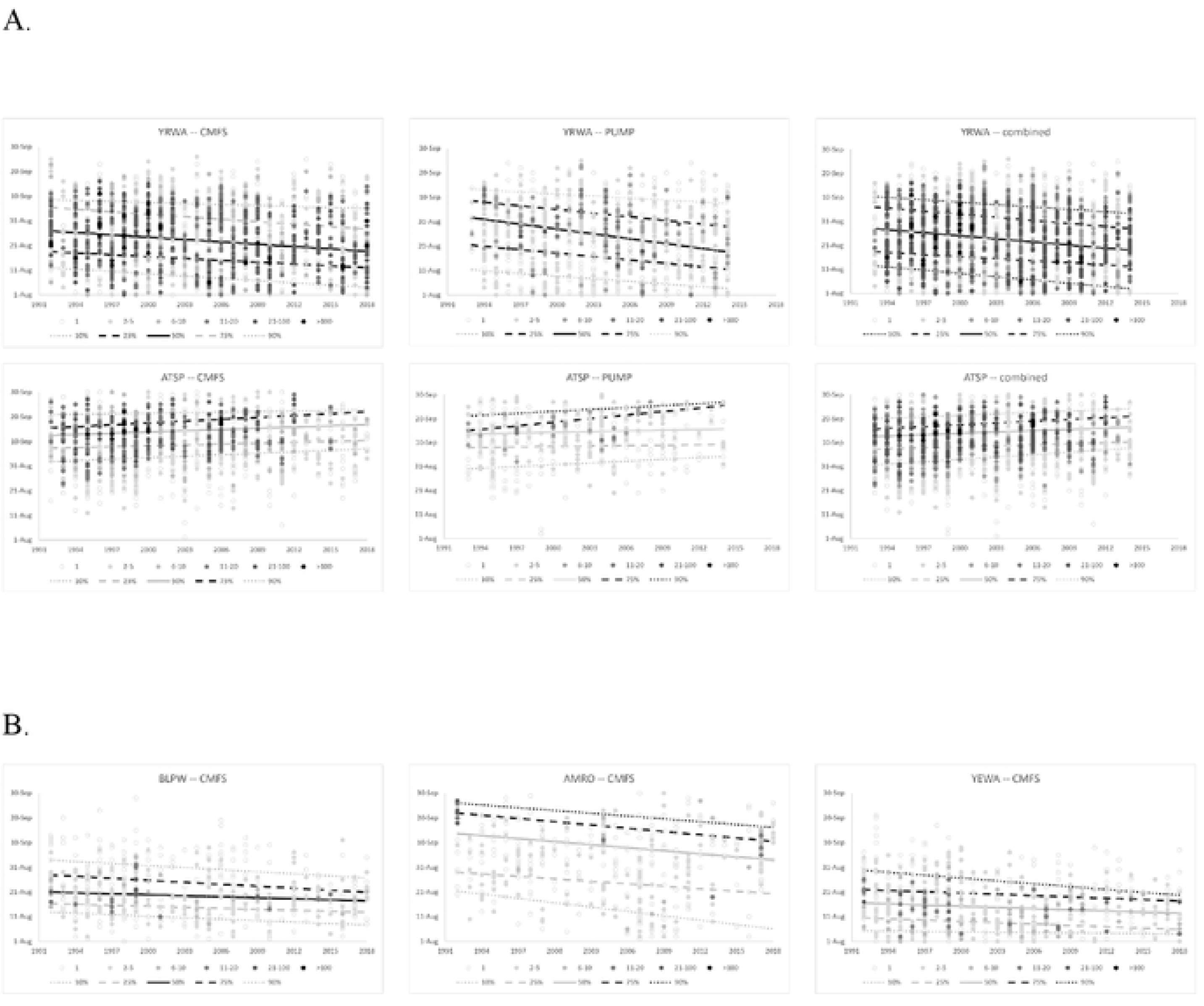
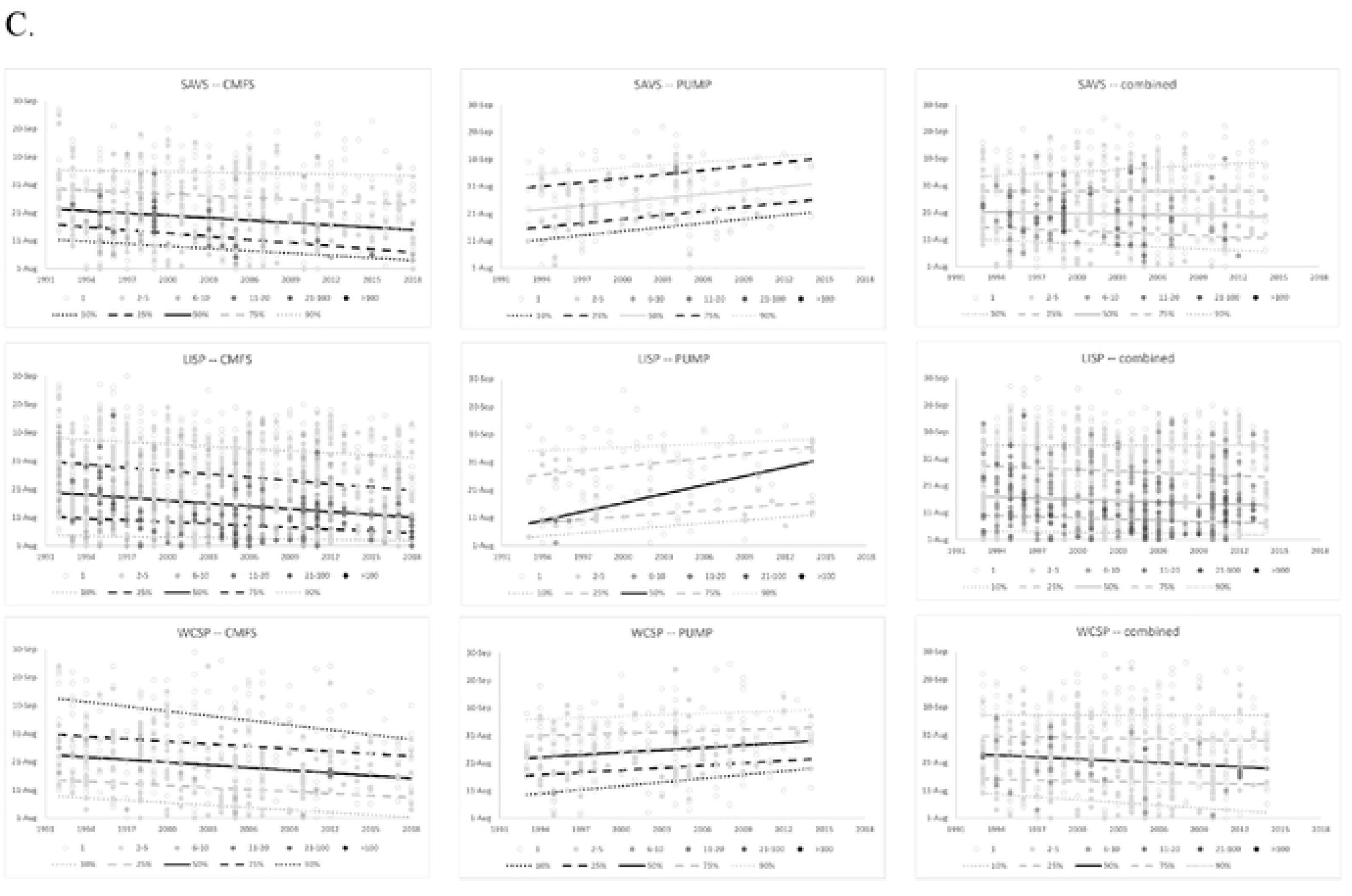
Quantile regression in the timing of autumn migration for select species of migratory species captured at Creamer’s Field Migration Station (CFMS site; Fairbanks, AK; 1992-2018) and the PUMP site (Tetlin, AK; 1993-2014). Both ages were combined for this quantile regression. Darker dots indicate more birds captured on that date. Dotted lines indicate 10^th^ and 90^th^ percentiles, dashed lines indicate 25^th^ and 75^th^ percentiles, with the solid line showing the median; black lines indicate when the 95% confidence interval of the slope did not include 0.

Results for all 21 species are shown in Supporting Information Figure S1 and Supporting Information Tables S4 and S5.

A. YRWA showed an advance in capture date over time and ATSP showed a delay at both sites.
B. Three examples (of 13 species) that showed evidence for advanced autumn migration timing only at the CFMS banding site.
C. Opposing, site-specific differences in timing of three sparrow species.

Interestingly, three sparrow species showed a consistent, site-specific pattern (Savannah Sparrow, Lincoln’s Sparrow, White-crowned Sparrow; Figure 5C), indicating advanced migration at CFMS but delayed migration at PUMP (Figure 5C). At CFMS, 13 additional species advanced in migratory timing, whereas only one was delayed (Figure 5A, B; Supporting Information Table S3). By contrast, at PUMP, three additional species advanced and 4 delayed (Figure 5A, Supporting Information Table S5).

Depending on species, capture location, and age group, the rate of advancement in median capture date per decade ranged between 1.3 days (Blackpoll Warbler) and 10.0 days (Yellow-rumped Warbler; Supporting Information Table S4), with species typically advancing by ∼2 to 3 days per decade (Supporting Information Tables S4, S5). Delays in median capture date ranged from 2.0 days (Dark-eyed Junco) to 11.1 days per decade (Lincoln’s Sparrow; Supporting Information Table S4). Long-distance migrants in our dataset frequently showed advanced migration timing with eight species at CFMS and two species at PUMP (Figure S1, Supporting Information Table S5). Short-distance migrants, however, lacked a consistent pattern (Supporting Information Table S5), due to between-site differences (Supporting Information Table S4), with five short-distance migrants advancing and one delaying at CFMS, and one short-distance migrant advancing and two delaying at PUMP.

Patterns of change in capture dates also differed when site data were combined. For some species (Swainson’s Thrush, Dark-eyed Junco, Gray-cheeked Thrush) there were no changes in capture dates at any quantile until both sites were combined (Supporting Information Tables S4, S5). For others, (Hermit Thrush, Yellow Warbler, Blackpoll Warbler, and Orange-crowned Warbler), site-specific trends became less evident (i.e., CI of the slope of quantile change included 0; Supporting Information Tables and Figure S4, S5, S1). There were also many instances where capture dates showed a time pattern at only one site, especially CFMS, which remained evident when both sites were combined. Yellow-rumped Warbler and American Tree Sparrow were the exceptions; Yellow-rumped Warbler exhibited strong, consistent patterns showing capture date advances for multiple quantiles, whether sites were treated separately or combined (Figure 5A, Supporting Information Tables S4, S5); a similar, consistent pattern was also seen with American Tree Sparrow but with capture delays rather than advances (Figure 5A, Supporting Information Tables S4, S5).

In some species, there were substantial differences in median capture date between age classes (Supporting Information Tables S4). Ten of twelve species of long-distance migrants showed differences between ages (CI did not overlap with 0) and eight of nine short-distance migrants. However, we caution that sample size may be a consideration here, as we captured many more HY birds each year than adults.

### Climate Indices and Avian Time Periods

For 12 species we found associations between long-term patterns of median capture dates (of both age classes combined) and adjusted climate indices (Supporting Information Figure S2, Supporting Information Table S7). Indices with the strongest patterns (i.e., most species with strong relationships) for each of the three time periods we defined are illustrated in Figure 6. The AO climate index during the Breeding Period was associated with the greatest number of species (9) that showed long-term changes in median capture dates (Figure 6, Supporting Table S7).

**Figure 6.**
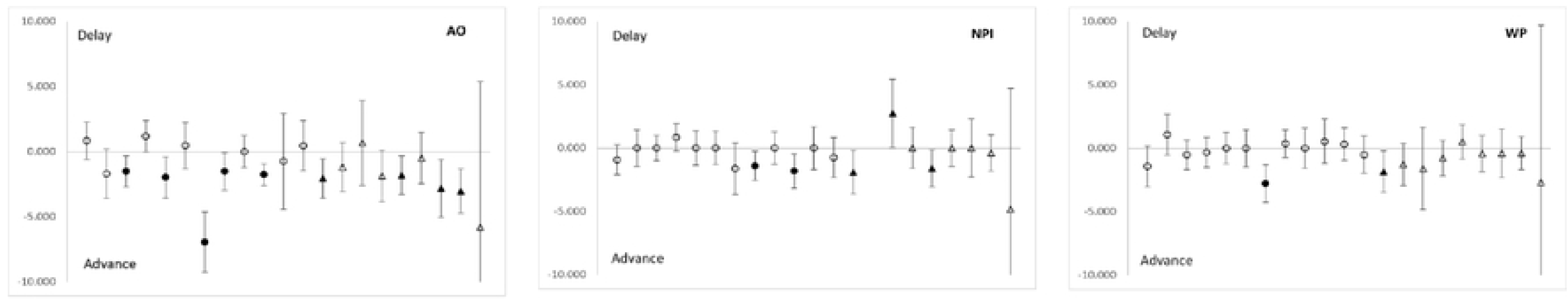
Example associations between period-specific climate indices (AO during Breeding, NPI during Post-fledging, and WP during Migration) and median capture dates for 21 bird species. During breeding, the AO index was consistently associated with advances in autumn migratory timing (species falling below the line), whereas there was less of a pattern for indices in the post-fledge and migration panels. Advances or delays in migratory timing are indicated for one standard deviation of change in the climate index. Change in capture timing (Y-axis) is reported as days change per decade with 85% confidence intervals. The first 12 species (left to right; circles) are long-distance migrants (ALFL, HAFL, GCTH, SWTH, OCWA, YEWA, YRWA, BLPW, NOWA, WIWA, SAVS, LISP) and the last 9 species (triangles) are short-distance migrants (RCKI, HETH, AMRO, VATH, ATSP, FOSP, WCSP, DEJU, RUBL). Closed circles and triangles are where the CI did not overlap 0. See Supporting Figure S2 for figures of all combinations of species, climate index, and period and Supporting Table S7 for parameter estimates and CI associated with the figures.

Decreasing index values of AO, which are associated with warmer temperatures (Bieniek et al. 2012, Bieniek et al. 2014), were predictive of an advanced median capture date (i.e., birds captured earlier in the autumn). Other indices were also associated with advanced capture dates, particularly during the Breeding Period (Supporting Information Figure S2, Table S7).

Compared to the Breeding Period, climate indices calculated during the Post-fledging and Migration Periods were less consistent in predicting a directional shift in median migration date among species (e.g., species fall both above and below the line in the right-hand panels of Figure 6; see also: Supporting Figure S2, Table S7). Predicted outcomes of some climate indices varied by time period as well. For example, for EPNP, warm springs (Breeding) led to advanced migration, while warm summer (Post-fledge) and autumn (Migration) periods were associated with delayed migration (Supporting Figure S2, Table S7). Finally, two climate indices, WP (during the Post-fledging Period) and NPI (during the Migration Period), did not predict any changes in median capture dates for any species.

## DISCUSSION

### Long-term changes in capture dates

Long-term advances in migration timing have occurred across all three North American flyways in recent decades, with the strongest patterns evident in spring, and fewer in autumn (e.g., Horton et al. 2020; reviewed in Gordo 2007). Our study focused on the autumn timing of 21 migratory passerines in boreal Alaska, a time of year and location that is generally understudied (Gallinat et al. 2015, Seavy et al. 2018), but an area that nonetheless exhibits steep avian declines and accelerated rates of climate change (e.g., Loarie et al. 2009, Rosenberg et al. 2019). As we predicted, we found evidence for advancement in at least one quantile of autumn passage in a majority (76%) of 21 species over a multi-decade time period, particularly among long-distance migrants captured at the CFMS study site (Figures 5, Supporting Figure S1, Supporting Tables S4, S5).

Though rates of long-term change are difficult to compare directly, advancement among passerines in boreal Alaska during autumn appears to be most common, compared to more variable outcomes in continental North America (Mills et al 2005, Ellwood et al. 2015, Barton and Sandercock 2018, Horton et al. 2020, 2023) and elsewhere (Van Buskirk et al. 2009; reviewed in Gordo 2007). Species in our study, for example, regularly advanced 2-3 days per decade in autumn (Figures 5, Supporting Figure S1, Supporting Tables S4, S5), compared to estimates of <1 day per decade in continental North America (Horton et al. 2020, 2023). We hypothesize that the accelerated rate of warming at high latitudes (e.g., Loarie et al. 2009, Ballinger et al. 2023) results in greater phenological shifts toward earlier breeding (e.g., Zaifman et al. 2017, Chmura et al. 2020, Horton et al. 2023; reviewed in Chmura et al. 2019). Earlier completion of breeding carries over to result in a pattern of earlier autumn departures (e.g., Chmura et al. 2020). Given that boreal Alaska exhibits a compressed breeding season (Benson and Winker 2015), where single-broods are the norm (Ringgenberg and Winker 2015), and many species rely on insect prey (e.g., flycatchers, warblers), the fitness benefits to individuals staying later in autumn may be minimal, compared to risks of cold end-of-season weather.

Long-distance migrants departing earlier in autumn are consistent with other investigations (e.g., Jenni and Kéry 2003, Van Buskirk et al. 2009, Gallinat et al. 2015), including data for Alaska (Zaifman et al. 2017). The distribution of capture dates at our boreal study sites also reflected an inherent pattern of earlier travel times for long-distance compared to short-distance species (Figure 3). A tendency for long-distance migrants to depart soon after breeding presumably reflects additional time required to travel greater distances (e.g., MacMynowski and Root 2007).

Although short-distance migrants often exhibit long-term delays in departure times (Jenni and Kéry 2003, Ellwood et al. 2015, Gallinat et al. 2015, Barton and Sandercock 2018), autumn data for boreal Alaska did not reveal a consistent pattern (Figures 5, Supporting Figure S1; Suporting Tables S4, S5). Benson and Winker (2015) also highlighted a substantial inter- specific variation in molt and fattening of autumn migrants following the brief breeding season in boreal Alaska. One short-distance species, however, stands out relative to others. American Tree Sparrow delayed timing across multiple portions (e.g., quantiles) of the autumn migratory period at both study sites (Figures 5, Supporting Figure S1). We hypothesize the pattern relates to one (or more) of the following mechanisms. First, the American Tree Sparrow is unusual relative to other boreal migrants, including other sparrows, in that it spends one of the longest durations on the breeding grounds, where nearly all complete prebasic molt, resulting in one of the lowest proportions of birds that exhibit molt during autumn migration (Benson and Winker 2015, Naugler et al. 2020). Molting in Alaska after breeding creates a temporal separation that presumably avoids an energy allocation conflict with migration (e.g., Tonra and Reudink 2018, Pageau et al. 2020). If climate change offers milder autumns with abundant, omnivorous feeding opportunities (e.g., seeds), then delayed departures over time may reflect a predisposition in this species to stay longer to molt and capitalize on fattening. Alternatively, long-term delays in American Tree Sparrow could also result from long-term cooling trends on the breeding grounds of the population(s) we sampled, causing delays in nesting that subsequently delayed autumn departures (see Site-Specific discussion, below; Halupka and Halupka 2017, Chmura et al. 2020).

### Site-specific and intra-specific patterns

Three sparrow species (Lincoln’s Sparrow, Savannah Sparrow, and White-crowned, showed identical, site-specific patterns in autumn timing, namely long-term advances at CFMS, but long-term delays at PUMP (Figure 5c). Differing site-specific trends highlights the need for caution before extrapolating migratory patterns over broad spatial scales (Mills 2005, Ellwood et al. 2015, Mayor et al. 2017). It is important to remember that the magnitude and direction of climate change vary in both space and time (e.g., Loarie et al. 2009, Knudsen et al. 2011, Mayor et al. 2017). Alaska’s immense size means that different areas of the state exhibit distinctive climate “regions” (Bieniek 2012), which can exhibit different rates of warming and cooling over time (e.g., Wendler et al. 2012, Bieniek et al. 2014, Mayor et al. 2017). This creates the potential for different populations of the same species to experience different environmental conditions during breeding.

Alaska’s immense size means that different areas of the state exhibit different and distinctive climate “regions” (Bieniek 2012), which show different long-term trends in warming and precipitation (Ballinger et al. 2023). Other environmental conditions also vary, such as the date marking the end of snow season (Ratanen et al. 2023, ACCAP 2023). Hence, depending on where a bird breeds in Alaska, the timing of spring green-up may differ (e.g., warmer sites with reduced snow exhibit earlier green-up than the reverse; Zheng et al. 2022). Spring green-up is an important predictor of avian breeding phenology, as it marks availability of insect food resources (e.g., Cole et al. 2015, Mayor et al. 2017).

We hypothesize that Alaska’s spatially-variable environmental conditions are one possible explanation for the opposing, site-specific trends in autumn migratory timing that we documented in three sparrow species (Figure 5c). If each banding site captured different breeding populations, those passing through CFMS may have experienced a warming trend on the breeding grounds, resulting in earlier spring green-up, nest completion, and initiation of migration. By contrast, birds at the PUMP site may have experienced the opposite scenario (cooler temperatures and/or delayed end of snow season over time), resulting in later nesting and a long-term delay in autumn captures. Our hypothesis not only applies to the intra-specific patterns of sparrows, but also to the broader site-specific pattern among taxa, in which species most often advanced autumn migratory timing at CFMS, but not at PUMP. Though we lack data on the exact breeding location(s) of autumn migrants caught at each station, future work may be able to coarsely infer (or exclude) candidate breeding regions in Alaska, based on predicted relationship(s) between the spatially-variable timing of green-up, onset of reproduction, and subsequent migratory passage.

Dunn (2005) emphasized the importance of analyzing banding sites separately, especially when it is unknown whether data represent a single population. Our two study sites were only 277 km apart, along the same migratory flyway (Tanana River valley), in comparable boreal habitats (Figure 1). Had we simply combined data as indicative of “central Alaska forests,” we might have overlooked important patterns because the site-specific trends (in opposing directions) would have negated each other (Figure 5c). Horton et al. (2023) recently combined autumn banding records that included the same two Alaska sites in a continent-wide analysis, but failed to detect long-term migratory patterns relative to temperature. One possible explanation is that climate at banding sites in autumn is a less important predictor of migration because it does not accurately reflect conditions during breeding, which our data suggest is closely associated with migratory timing.

We hesitate to infer any conclusions about long-term patterns in migratory timing by age-class (Supporting Table S4), as our results were likely influenced by uneven samples of many more hatch year birds captured relative to adults. However, two age-related outcomes are worth noting. First, hatch year birds typically migrated earlier in autumn than adults (Supporting Table S3), which presumably reflects the need for adults to replace flight feathers before departure (Benson and Winker 2001; reviewed in Newton 2011). Second, Alder Flycatchers were the exception in that adults departed earlier, a pattern presumably related to the adult ability to suspend post-breeding molt following one of the shortest breeding periods of any boreal passerine (Benson and Winker 2001).

### Climate Indices and Avian Time Periods

Our exploration of climate indices as predictors of median autumn capture date resulted in several important outcomes. First, climate indices indicative of warm Breeding Periods, rather than other time periods (e.g., Post-Fledging or Migration), were often associated with earlier autumn migration (Figure 6, Supporting Figure S2, Supporting Table S7). Second, warming associated with the AO climate index, in particular, was associated with advanced migration in the greatest number of species in our dataset (9 or 43% of taxa; Figure 6, Supporting Table S7).

Collectively, our results support the hypothesis that climate conditions, particularly those associated with warming on the breeding grounds, are likely to produce earlier autumn passage of Alaska’s boreal migrants. This is perhaps not surprising, as temperatures are the ultimate driver of migration (Gordo 2007). In boreal Alaska, shifts in atmospheric circulation have increased temperatures and decreased snowfall, advancing spring snowmelt by eight days since the mid-1960s (Euskirchen et al. 2009). Given generally warmer, localized conditions in spring, birds can complete breeding earlier and therefore depart sooner (Zaifman 2017, Chmura et al. 2020). Accumulated evidence outside Alaska also supports the principle that breeding phenology influences the timing of autumn migration (Mitchell et al. 2012; Briedis et al. 2018; Fayet et al. 2016; Saino et al. 2017; Gow et al. 2019; De Zwaan et al. 2019, Chmura et al. 2020).

The AO index, a predictor of advanced autumn timing of median migration date in our dataset, is a leading mode of sea level pressure variability in the Northern Hemisphere, with strong wintertime activity that persists through summer and into fall (Thompson and Wallace 1998, Bond and Harrison 2006, Wang et al. 2018). AO’s climatic trend toward warmer temperatures has been associated with thinning sea ice in the North Pacific since 1979, ultimately affecting warming patterns at high latitudes (Rinke et al. 2013, Vihma 2014, Wang et al. 2019). Yet, outside of Alaska, studies of the North Atlantic Oscillation (NAO), a climate pattern that is highly correlated with the annular structure of AO (Gong et al. 2018), yielded surprisingly little association with spring or autumn capture dates (Marra et al. 2005, Wilson 2007, MacMynowski and Root 2007, Miller-Rushing et al. 2008, DeLeon et al. 2011, and Barton and Sandercock 2018; *cf.* Van Buskirk 2009). This suggests that perhaps AO (or other climate indices like it) are more relevant to boreal Alaska. It seems plausible that the AO index (or others) serves as a proxy for the timing of spring green-up and other mechanistic drivers(s) of migratory patterns, which will require further field study to reveal. Further studies are also needed to differentiate the relationship between climate indices or other indices of change in climate such as ERA-5 Land reanalysis data set (Balsamo et al. 2015, Ratanen et al. 2023) and long-term changes in the timing of migration.

We caution that median migration date may not be the best response metric to understand climate relationships, even though it has been used in other investigations (e.g., Barton and Sandercock 2018). The leading edge (early migrants) or trailing edge (late migrants) of migratory passage are potentially more sensitive to change (MacMynowski and Root 2007), as evidenced by studies (including our own) that report migratory patterns based on quantiles (Tøttrup et al. 2006, Wilson 2007, Miller-Rushing et al. 2008, Barton and Sandercock 2018).

Further analysis is needed to quantify whether climate indices such as AO (or others) are more tightly correlated with other quantiles, or show any site-specific differences, given spatially distinctive climate conditions in Alaska (Bienick et al. 2012, 2014; Wendler 2012).

## CONCLUSIONS AND CONSERVATION IMPLICATIONS

Long-term changes, especially advances, in autumn migration are occurring in boreal Alaska’s passerines, and these are associated with a warming climate during breeding months. However, we advise caution before extrapolating this pattern across broad spatial extents in Alaska, as spatial heterogeneity in regional climate may differentially impact breeding populations, resulting in striking site-specific and species-by-site patterns. Understanding autumn migration in the context of climate, including statewide spatial patterns of advancing or delayed spring green-up, gives managers of migratory birds new insight into how potential stressors may affect different breeding populations across Alaska.

Ultimately, a bird’s ability to cope with a rapidly changing environment depends upon its rate of adaptive response (Knudsen et al. 2011), but there appears to be a general lag in the ability of avian species to keep up (Radchuk et al. 2019). Banding stations often provide the only source of data on bird population parameters (e.g., Ralph et al. 1993, Dunn 2005, Desante et al. 2018), while also providing opportunities to quantify mechanisms that birds use to cope with climate change, which can be strongly species-specific (Carey 2009). Examples of avian coping mechanisms include altered timing of annual cycles (e.g., Burrows et al. 2011, Jukema et al. 2014, Mayor et al. 2017), the extent of molt (Kiat et al. 2019), and changes in body size (e.g. Van Buskirk et al. 2010, Jirinec et al. 2021, Radchuk et al. 2019) or proportion (Goodman et al. 2012). These metrics are typically contained in banding records.

CFMS in Fairbanks is currently the only remaining long-term avian banding station in Alaska. This uniquely positions its valuable data resources for future investigations of coping mechanisms (or lack thereof) that boreal birds exhibit at high latitudes. For example: Are species that show little evidence for advances in migratory timing more likely to show population declines? (Møller et al. 2008; Iler et al. 2021). However, quantification of CMFS data alone is less effective than a network of banding stations in Alaska. This would allow us to anticipate, over broader spatial scales, which species (or populations) are most vulnerable to Alaska’s rapidly changing environment and implement effective conservation strategies (Urban et al. 2015).

## ACKNOWLEDGEMENTS

Thank you to the anonymous reviewers for providing feedback on this article, and also to JJ Frost, Robert Snowden, Chris Barger, and Tracy Gotthardt for the edits before submission. A huge thank you to all the past and present staff, volunteers, board members, students, interns, community members, Adopt-A-Net Sponsors, and collaborators for spending thousands of hours pouring their love and expertise into both CFMS and PUMP. And to the Alaska Bird Observatory (1992-2012), who provided much of the data from CFMS. The authors would also like to thank Dan Gibson, Kevin Winker, Nancy Fresco, Rick Thoman, and the Klamath Bird Observatory for their advice and expertise.

## SUPPORTING INFORMATION

**Table S1. Total numbers of individuals and proportional demographic composition by age and sex for 21 passerine species at Creamer’s Field Migration Station (CFMS, Fairbanks, AK) and the Pump Station (PUMP, near Tok, AK); capture years were 1992-2018 for CFMS and 1993-2014 for PUMP**.

**Table S2. Species-specific Intervals used for calculating climate indices used as predictor variables in regression analyses of capture dates,** (see also Figure 2)

**Table S3. Median capture dates (95% CI) used to model long-term changes in the migration timing of 21 passerine species at Creamer’s Field Migration Station (CFMS in Fairbanks, AK; 1992-2018) and the Pump Station ((PUMP near Tok, AK; 1993-2014).** Differences in median capture dates (in days) between ages and between sites for each species are also given. Bold indicates that the CI of the difference between medians did not overlap in 0.

**Table S4. Estimated changes (95% CI) in median capture dates by site and age.** Estimates are predicted number of days in change of the median capture date per decade. Bold indicates 95% CI did not include 0.

**Table S5. Estimated changes (95% CI) of quantiles of capture distributions for combined ages and site data (CFMS and PUMP); estimates are days of changes per decade.** Bold entries indicate estimates whose 95% CI do not include 0. Estimates are shaded to indicate species and quantiles where we found differences in slopes between CFMS and PUMP (separate estimates not included in the table, but see Supplemental Fig. 1), suggesting the need for caution in interpretation.

**Table S6. Estimated change in the median capture date for one unit of change in each adjusted climate index and time period with confidence intervals.**

**Table S7: Estimated change in the median capture date for one standard deviation of change in each adjusted climate index and time period with confidence intervals** (see also Figure 6, Supplemental S2 Figure).

**Figure S1. Quantile regression of the timing of autumn migration (day 274 = October 1) for 21 passerine species at Creamer’s Field Migration Station (Fairbanks, AK) and the PUMP site (Tetlin, AK); capture years were 1992-2018 for CFMS and 1993-2014 for PUMP.** Darker dots indicate more birds captured on that date. Dotted lines indicate 10^th^ and 90^th^ percentiles, dashed lines indicate 25^th^ and 75^th^ percentiles, with the solid line showing the median; black lines indicate when the 95% confidence interval of the estimated change did not include 0.

**Figure S2: Estimated changes in median capture date vs. climate indices.** Changes in capture timing are reported in days per SD of climate index with 85% confidence intervals. For ease of comparison, NPI and AO (higher index = colder) figures are based on a 1 SD decline in the index, while all other indices (higher index = warmer) are based on a 1 SD increase in the index. The first 12 species (left to right (circles) are long-distance migrants (ALFL, HAFL, GCTH SWTH, OCWA, Y, YRWA, BLPW, NOWA, WIWA, SAVS, LISP) and the last 9 species (triangles) are short-distance migrants (RCKI, HETH, AMRO, VATH, ATSP, FOSP, WCSP, DEJU, RUBL). The left column of figures is for nesting/breeding period calculated indices, the middle column is for predictors calculated for the post-fledging to mid-migration period, and the right column is for predictors calculated for the migration period.

## Notes

### Competing Interest Statement

The authors have declared no competing interest.

## REFERENCES

ACCAP. 2023. Alaska Center for Climate Assessment and Policy. “Change in end of snow season (1950–2021)” graphic. https://uaf-accap.org/accap-projects/graphics/. Accessed on 15 May 2023.

ADFG (2015). Alaska Wildlife Action Plan. Alaska Department of Fish and Game. Juneau, AK, USA. http://www.adfg.alaska.gov/index.cfm?adfg=species.wapview

Anders, A. D., J. Faaborg, and F. R. Thompson III (1998). Postfledging dispersal, habitat use, and home-range size of juvenile Wood Thrushes. The Auk 115: 349–358.

Ballinger, T. J., U. S. Bhatt, P. A. Bieniek, B. Brettschneider, R. T. Lader, J. S. Littell, R. L. Thoman, C. F. Waigl, J. E. Walsh, and M. A. Webster (2023). Alaska terrestrial and marine climate trends, 1957-2021. Journal of Climate 10:1–41.

Balsamo, G., Albergel, C., Beljaars, A., Boussetta, S., Brun, E., Cloke, H., Dee, D., Dutra, E., Muñoz-Sabater, J., Pappenberger, F., de Rosnay, P., Stockdale, T., and Vitart, F. (2015): ERA-Interim/Land: a global land surface reanalysis data set, Hydrology and Earth System Sciences 19: 389–407, 10.5194/hess-19-389-2015.

Bateman, B.L, C. Wilsey, L. Taylor, J. Wu, G.S. LeBaron, G. Langham (2020). North American birds require mitigation and adaptation to reduce vulnerability to climate change. Conservation Science and Practice 2: 8. https://conbio.onlinelibrary.wiley.com/doi/full/10.1111/csp2.242

Barnston, A. G., and R. E. Livezey (1987). Classification, seasonality and persistence of low- frequency atmospheric circulation patterns. Monthly weather review 115: 1083–1126.

Barton, G. G., and B. K. Sandercock (2018). Long-term changes in the seasonal timing of landbird migration on the Pacific flyway. Ornithological Applications 120:30–46.

Baxter, S., and S. Nigam (2015). Key role of the North Pacific Oscillation–west Pacific pattern in generating the extreme 2013/14 North American winter. Journal of Climate 28: 8109–8117.

Benson, A. M., and K. Winker (2015). High-latitude passerine migrants overlap energetically demanding events in autumn. The Wilson Journal of Ornithology 127: 601–614.

Benson, A. M., W. N. Johnson, R. P. Barry, and S. L. Guers (2012). Evaluation of autumn mist-netting data for monitoring passerine populations in interior Alaska. Wildlife Society Bulletin 36: 328–335.

Benson, A. M., and K. Winker (2001). Timing of breeding range occupancy among high-latitude passerine migrants. The Auk., 118:513–9.

Bieniek, P. A., U. S. Bhatt, L. A. Rundquist, S. D. Lindsey, X. Zhang, and R. L. Thoman (2011). Large-scale climate controls of interior Alaska river ice breakup. Journal of Climate 24: 286–297.

Bieniek, P. A., U. S. Bhatt, R. L. Thoman, H. Angeloff, J. Partain, J. Papineau, J., F. Fritsch, E. Holloway, J. E. Walsh, C. Daly, M. Shulski, et al. (2012). Climate divisions for Alaska based on objective methods. Journal of Applied Meteorology and Climatology 51: 1276–1289.

Bieniek, P. A., J. E. Walsh, R. L. Thoman, and U. S. Bhatt (2014). Using climate divisions to analyze variations and trends in Alaska temperature and precipitation. Journal of Climate 27: 2800–2818.

Billerman S. M., B. K. Keeney, P. G. Rodewald, and T. S. Schulenberg (Editors) (2022). Birds of the World. Cornell Laboratory of Ornithology, Ithaca, NY, USA.

BirdLife International (2023) Important Bird Areas factsheet: Upper Tanana River Valley. Downloaded from http://www.birdlife.org on 12/04/2023. http://datazone.birdlife.org/site/factsheet/upper-tanana-river-valley-iba-usa https://birdsoftheworld.org/bow/home

Bond, N. A., and D. E. Harrison (2006). ENSO’s effect on Alaska during opposite phases of the Arctic Oscillation. International Journal of Climatology: A Journal of the Royal Meteorological Society 26: 1821–1841.

Briedis, M., M. Krist, M. Král, C. C. Voigt, and P. Adamík (2018). Linking events throughout the annual cycle in a migratory bird—non-breeding period buffers accumulation of carry- over effects. Behavioral Ecology and Sociobiology 72: 1–12.

Brown, J. M., and P. D. Taylor (2015). Adult and hatch-year blackpoll warblers exhibit radically different regional-scale movements during post-fledging dispersal. Biology Letters 11: 20150593.

Bureau of Land Management (2019). Bureau of Land Management Alaska Special Status Species List. [Accessed 21 March 2022] https://www.blm.gov/sites/blm.gov/files/uploads/Alaska_Special-Status-Species-List_2019.pdf

Burrows, M. T., D. S. Schoeman, L. B. Buckley, P. Moore, E. S. Poloczanska, K. M. Brander, C. Brown, J. F. Bruno, C. M. Duarte, B. S. Halpern, J. Holding, et al. (2011). The pace of shifting climate in marine and terrestrial ecosystems. Science 334: 652–655.

Cade, B. S., and B. R. Noon (2003). A gentle introduction to quantile regression for ecologists. Frontiers in Ecology and the Environment 1: 412–420.

Carey, C. (2009). The impacts of climate change on the annual cycles of birds. Philosophical Transactions of the Royal Society B: Biological Sciences, 364: 3321–3330.

Chmura, H. E., Kharouba, H. M., Ashander, J., Ehlman, S. M., Rivest, E. B. and L. H. Yang (2019). The mechanisms of phenology: the patterns and processes of phenological shifts. Ecological monographs, 89: p.e01337.

Chmura, H. E., J. S. Krause, J. H. Pérez, M. Ramenofsky, and J. C. Wingfield (2020). Autumn migratory departure is influenced by reproductive timing and weather in an Arctic passerine. Journal of Ornithology 161:779–791.

Cole, E.F., Long, P.R., Zelazowski, P., Szulkin, M. and Sheldon, B.C., 2015. Predicting bird phenology from space: Satellite-derived vegetation green-up signal uncovers spatial variation in phenological synchrony between birds and their environment. Ecology and Evolution, 5: 5057–5074. 10.1002/ece3.1745

Cooper, B. A., and R. J. Ritchie (1995). The Altitude of Bird Migration in East-Central Alaska: A Radar and Visual Study. Journal of Field Ornithology 66: 590–608. http://www.jstor.org/stable/4514063

DeLeon, R. L., E. E. DeLeon, and G. R. Rising (2011). Influence of climate change on avian migrants’ first arrival dates. The Condor 113: 915–923.

DeSante, D. F., Kaschube, D. R., and Saracco, J. F. 2018. Population changes and their demographic drivers in landbirds of western North America: An assessment from the Monitoring Avian Productivity and Survivorship program, in Trends and traditions: Avifaunal change in western North America (W. D. Shuford, R. E. Gill Jr., and C. M. Handel, eds.), pp. 269–293. Studies of Western Birds 3. Western Field Ornithologists, Camarillo, CA; doi 10.21199/SWB3.15.

Dunn, E. H. (2005). Counting migrants to monitor bird populations: state of the art. USDA Forest Service General Technical Report PSW-GTR-191.

Department of Defense Partners in Flight (2021). Department of Defense Mission Sensitive Species. https://denix.osd.mil/dodpif/mss-featured-content/products/mss-factsheet/. Accessed on [21 March 2022].

Douglas, C.A., and C. Zhang (2021). Machine learning analyses of remote sensing measurements establish strong relationships between vegetation and snow depth in the boreal forest of Interior Alaska. Environmental Research Letters 16:065014.

De Zwaan, D. R., S. Wilson, E. A. Gow, and K. Martin (2019). Sex-specific spatiotemporal variation and carry-over effects in a migratory alpine songbird. Frontiers in Ecology and Evolution 7, 285.

Ellwood, E. R., A. Gallinat, R. B. Primack, and T. L. Lloyd-Evans (2015). Autumn migration of North American landbirds. Pp. 193–205 in E. M. Wood, and J. L. Kellermann (Editors), Phenological synchrony and bird migration: changing climate and seasonal resources in North America. Studies in Avian Biology (no. 47), CRC Press, Boca Raton, FL.

Euskirchen, E. S., A. D. McGuire, F. S. Chapin III, S. Yi, and C. C. Thompson (2009). Changes in vegetation in northern Alaska under scenarios of climate change, 2003–2100: implications for climate feedbacks. Ecological applications 19: 1022–1043.

Fair, J., E. Paul, J. Jones, and L. Bies, Eds. 2023. Guidelines to the Use of Wild Birds in Research. Ornithological Council. http://www.BIRDNET.org.

Fayet, A. L., R. Freeman, A. Shoji, H. L. Kirk, O. Padget, C. M. Perrins, and T. Guilford (2016). Carry-over effects on the annual cycle of a migratory seabird: an experimental study. Journal of animal ecology 85: 1516–1527.

Fink, D., T. Auer, A. Johnston, M. Strimas-Mackey, O. Robinson, S. Ligocki, W. Hochachka, L. Jaromczyk, C. Wood, I. Davies, M. Iliff, and L. Seitz (2021). eBird Status and Trends, Data Version: 2020; Released: 2021. Cornell Lab of Ornithology, Ithaca, NY, USA. 10.2173/ebirdst.2020

Gallinat, A. S., R. B. Primack, and D. L. Wagner (2015). Autumn, the neglected season in climate change research. Trends in Ecology & Evolution 30(3), 169–176. 10.1016/j.tree.2015.01.004

Gibson, D. (2011). Nesting shorebirds and landbirds of Interior Alaska. Report for US Geological Survey, Alaska Science Center, Anchorage.

Gong, H., L. Wang, W. Chen, and D. Nath (2018). Multidecadal fluctuation of the wintertime Arctic Oscillation pattern and its implication. Journal of Climate 31: 5595–5608.

Goodman, R.E., Lebuhn, G., Seavy, N.E., Gardali, T. and Bluso-Demers, J.D. (2012), Avian body size changes and climate change: warming or increasing variability?. Glob Change Biol, 18: 63–73. 10.1111/j.1365-2486.2011.02538.x

Gordo, O., and J. J. Sanz (2006). Climate change and bird phenology: a long-term study in the Iberian Peninsula. Global Change Biology 12: 1993–2004.

Gordo, O. (2007). Why are bird migration dates shifting? A review of weather and climate effects on avian migratory phenology. Climate research 35: 37–58.

Gow, E. A., L. Burke, D. W. Winkler, S. M. Knight, D. W. Bradley, R. G. Clark, M. Bélisle, L. L. Berzins, T. Blake, E. S. Bridge, R. D. Dawson at al. (2019). A range-wide domino effect and resetting of the annual cycle in a migratory songbird. Proceedings of the Royal Society B 286(1894), 20181916.

Halupka, L., and K. Halupka (2017). The effect of climate change on the duration of avian breeding seasons: a meta-analysis. Proceedings of the Royal Society B: Biological Sciences, 284(1867), 20171710. 10.1098/rspb.2017.1710

Handel, C.M., I. J. Stenhouse, and S. M. Matsuoka (Editors) (2021). Alaska Landbird Conservation Plan, version 2.0. Boreal Partners in Flight, Anchorage, AK. 146 pp.

Handel, C. M., and J. R. Sauer (2017). Combined analysis of roadside and off-road breeding bird survey data to assess population change in Alaska. The Condor: Ornithological Applications 119.3: 557–575.

Harding Scurr, A., J, Hagelin, G. Pendleton, K. DuBour, T. Blake, C. Stuyck, E. Allaby. Data from: Long-term changes in the timing of autumn migration in Alaska’s boreal songbirds. Ornithology / Ornithological Applications. Data DOI or URL.

Hartmann, B., and G. Wendler (2005). The significance of the 1976 Pacific climate shift in the climatology of Alaska. Journal of climate 18: 4824–4839.

Horton, Kyle, Morris, Sara, Van Doren, Benjamin, Covino, Kristen. 2023. Six decades of North American bird banding records reveal plasticity in migration phenology. Journal of Animal Ecology. 92: 738–750

Horton, K. G., F. A. La Sorte, D. Sheldon, T. Y. Lin, K. Winner, G. Bernstein, S. Maji, W. M. Hochachka, and A. Farnsworth (2020). Phenology of nocturnal avian migration has shifted at the continental scale. Nature Climate Change 10: 63–68.

Hussell, D. J. T., F. Bairlein, and E. H. Dunn (2014). Double brooding by the Northern Wheatear on Baffin Island. Arctic 67:167–172.

Iler, A. M., P. J. CaraDonna, J. R. Forrest, and E. Post (2021). Demographic consequences of phenological shifts in response to climate change. Annual Review of Ecology, Evolution, and Systematics 52: 221–245.

IUCN (2021). The IUCN Red List of Threatened Species. Version 2021–3. https://www.iucnredlist.org. Accessed on [21 March 2022].

Jenni, L., and M. Kéry (2003). Timing of autumn bird migration under climate change: advances in long–distance migrants, delays in short–distance migrants. Proceedings of the Royal Society of London. Series B: Biological Sciences 270: 1467–1471.

Jirinec V, Burner RC, Amaral BR, Bierregaard RO Jr, Fernández-Arellano G, Hernández-Palma A, Johnson EI, Lovejoy TE, Powell LL, Rutt CL, Wolfe JD, Stouffer PC. Morphological consequences of climate change for resident birds in intact Amazonian rainforest. Sci Adv. 2021 Nov 12;7(46):eabk1743. doi: 10.1126/sciadv.abk1743. Epub 2021 Nov 12. PMID: 34767440; PMCID: PMC8589309.

Jukema, J. & Wiersma, P. (2014). Climate change and advanced primary moult in Eurasian golden plovers pluvialis apricaria. Ardea 102: 153–160.

Kiat, Y., Vortman, Y. & Sapir, N. (2019). Feather moult and bird appearance are correlated with global warming over the last 200 years. Nat Commun 10, 2540. 10.1038/s41467-019-10452-1

Kessel, B., and D. D. Gibson (1978). Status and Distribution of Alaska Birds. Studies in Avian Biology, No. 1. Cooper Ornithological Society, Los Angeles, CA.

Kessel, B. 1984. Migration of sandhill cranes, Grus canadensis, in east-central Alaska, with routes through Alaska and western Canada. Canadian Field-Naturalist 98:279–282.

Knudsen, E., A. Lindén, C. Both, N. Jonzén, F. Pulido, N. Saino, W. Sutherland, L. A. Bach, T. Coppack, T. Ergon, P. Gienapp, and N. C. Stenseth (2011). Challenging claims in the study of migratory birds and climate change. Biological Reviews 86: 928–946.

Knudsen, E., A. Lindén, T. Ergon, N. Jonzén, J. O. Vik, J. Knape, J. E. Røer, and N. C. Stenseth (2007). Characterizing bird migration phenology using data from standardized monitoring at bird observatories. Climate Research 35: 59–77.

L’Heureux, M. L., M. E. Mann, B. I. Cook, B. E. Gleason, and R. S. Vose (2004). Atmospheric circulation influences on seasonal precipitation patterns in Alaska during the latter 20th century. Journal of Geophysical Research: Atmospheres, 109(D6).

Lindén, A. (2018). Adaptive and nonadaptive changes in phenological synchrony. Proceedings of the National Academy of Sciences, 115: 5057–5059.

Loarie, S., P. Duffy, H. Hamilton, G. P. Asner, C. B. Field, and D. D. Ackerly (2009). The velocity of climate change. Nature 462:1052–55. 10.1038/nature08649

Loss, S. R., B. V. Li, L. C. Horn, M. R. Mesure, L. Zhu, T. G. Brys, A. M. Dokter, J. A. Elmore, R. E. Gibbons, T. Z. Homayoun, K. G. Horton, et. al (2023). Citizen science to address the global issue of bird–window collisions. Frontiers in Ecology and the Environment. 10.1002/fee.2614

MacMynowski, D. P., and T. L. Root (2007). Climate and the complexity of migratory phenology: sexes, migratory distance, and arrival distributions. International Journal of Biometeorology 51: 361–373.

Malone, T., J. Liang, and E. C. Packee (2009). Cooperative Alaska forest inventory (Vol. 785). US Department of Agriculture, Forest Service, Pacific Northwest Research Station.

Markon, C., S. Gray, M. Berman, L. Eerkes-Medrano, T. Hennessy, H. Huntington, J. Littell, M. McCammon, R. Thoman, and S. Trainor (2018). Alaska. In Impacts, Risks, and Adaptation in the United States: Fourth National Climate Assessment, Volume II (D. R. Reidmiller, C. W. Avery, D. R. Easterling, K. E. Kunkel, K. L. M. Lewis, T. K. Maycock, and B.C. Stewart, Editors). U.S. Global Change Research Program, Washington, DC, USA 1185–1241. doi: 10.7930/NCA4.2018.CH26

Marra, P. P., C. M. Francis, R. S. Mulvihill, and F. R. Moore (2005). The influence of climate on the timing and rate of spring bird migration. Oecologia, 142: 307–315.

Matsuoka, S., J. Hagelin, M. Smith, T. Paragi, A. Sesser, and M. Ingle (2019). Pathways for avian science, conservation, and management in boreal Alaska. Avian Conservation and Ecology, 14(1).

Mayor, S. J., R. P. Guralnick, M. W. Tingley, J. Otegui, J. C. Withey, S. C. Elmendorf, M. E. Andrew, S. Leyk, I. S. Pearse, and D. C. Schneider (2017). Increasing phenological asynchrony between spring green-up and arrival of migratory birds. Scientific reports 7(1), 1–10.

McIntyre, C. L., and R. E. Ambrose. 1999. Raptor migration in autumn through the Upper Tanana River Valley, Alaska. Western Birds 30:33–38.

Miller-Rushing, A. J., T. L. Lloyd-Evans, R. B. Primack, and P. Satzinger (2008). Bird migration times, climate change, and changing population sizes. Global Change Biology 14: 1959–1972.

Mills, A. M. (2005). Changes in the timing of spring and autumn migration in North American migrant passerines during a period of global warming. Ibis 147: 259–269.

Mitchell, G. W., A. E. Newman, M. Wikelski, and D. Ryan Norris (2012). Timing of breeding carries over to influence migratory departure in a songbird: an automated radiotracking study. Journal of Animal Ecology 81:1024–1033.

Mitchell, G. W., P. D. Taylor, P. D, and I. G. Warkentin (2010). Assessing the function of broad- scale movements made by juvenile songbirds prior to migration. The Condor 112: 644–654.

Mizel, J. D., J. H. Schmidt, C. L. McIntyre, and M. S. Lindberg (2017). Subarctic-breeding passerines exhibit phenological resilience to extreme spring conditions. Ecosphere 8(2):e01680.

Møller, A. P., Rubolini, D., & Lehikoinen, E. (2008). Populations of migratory bird species that did not show a phenological response to climate change are declining. Proceedings of the National Academy of Sciences 105: 16195–16200.

Morton, M. L., M. W. Wakamatsu, M. E. Pereyra, and G. A. Morton (1991). Postfledging dispersal, habitat imprinting, and philopatry in a montane, migratory sparrow. Ornis Scandinavica 1: 98–106.

Morton, M. L. (1991). Postfledging dispersal of Green-tailed Towhees to a subalpine meadow. Condor 466–468.

Naugler, C. T., P. Pyle, and M. A. Patten, (2020). American Tree Sparrow (*Spizelloides arborea*), version 1.0. Birds of the World. Ithaca (NY): Cornell Lab of Ornithology. 10.2173/bow.amtspa.01

Newton, I. (2011). Migration within the annual cycle: species, sex and age differences. Journal of Ornithology 152: 169–185.

Pageau, C., Tonra, C. M., Shaikh, M., Flood, N. J., & Reudink, M. W. (2020). Evolution of moult-migration is directly linked to aridity of the breeding grounds in North American passerines. Biology Letters 16: 20200155.

Papineau, J. M. (2001). Wintertime temperature anomalies in Alaska correlated with ENSO and PDO. International Journal of Climatology: A Journal of the Royal Meteorological Society 21: 1577–1592.

Panjabi, A. O., W. E. Easton, P. J. Blancher, A. E. Shaw, B. A. Andres, C. J. Beardmore, A. F. Camfield, D. W. Demarest, R. Dettmers, R. H. Keller, K. V. Rosenberg, et al. (2020). Avian Conservation Assessment Database Handbook, version 2020. Partners in Flight Technical Series no. 8.1. 69 pp. https://pif.birdconservancy.org/wp-content/uploads/2021/02/acad.handbook.pdf

Perlwitz, J., T. Knutson, J. P. Kossin, and A. N. LeGrande (2017). Large-scale circulation and climate variability. In: Climate Science Special Report: Fourth National Climate Assessment, Volume I (D. J. Wuebbles, D. W. Fahey, K. A. Hibbard, D. J. Dokken, B. C. Stewart, and T. K. Maycock, Editors). U.S. Global Change Research Program, Washington, DC, USA, pp. 161–184. doi:10.7930/J0RV0KVQ

Pyle, P. 2022. Identification Guide to North American Birds, Part 1, 2nd Edition. Slate Creek Press, Forest Knolls, California. 698 pp.

Pyle, P. (1997). Identification guide to North American birds: a compendium of information on identifying, ageing, and sexing “near-passerines” and passerines in the hand. Slate Creek Press.

Radchuk, V., Reed, T., Teplitsky, C., van de Pol, M., Charmantier, A., Hassall, C., … & Kramer-Schadt, S. (2019). Adaptive responses of animals to climate change are most likely insufficient. Nature communications, 10: 3109.

Ralph, C. J. (1993). Handbook of field methods for monitoring landbirds (Vol. 144). Pacific Southwest Research Station.

Ralph, C.J.; Geupel, G.R.; Pyle, P.; Martin, T.E.; DeSante, D.F. 1993. Handbook of field methods for monitoring landbirds. Gen. Tech. Rep. PSW-GTR-144. Albany, CA: U.S. Department of Agriculture, Forest Service, Pacific Southwest Research Station. 41 p.

Ramenofsky, M., J. M. Cornelius, and B. Helm (2012). Physiological and behavioral responses of migrants to environmental cues. Journal of Ornithology 153: 181–191.

Ringgenberg, B., and K. Winker (2015). Indications that the Common Redpoll is double brooded in Alaska. Western Birds 46: 291–298.

Rinke, A., K. Dethloff, W. Dorn, D. Handorf, and J. C. Moore (2013). Simulated Arctic atmospheric feedbacks associated with late summer sea ice anomalies. Journal of Geophysical Research: Atmospheres 118: 7698–7714.

Rosenberg, K. V., M. A. Dokter, J. P. Blancher, J. R. Sauer, A. C. Smith, P. A. Smith, J. C. Stanton, A. Panjabi, L. Helft, M. Parr, and P. P. Marra (2019). Decline of the North American avifauna. Science, 366: 120–124. doi: 10.1126/science.aaw1313

Royama, T. (1992). Analytical population dynamics. Chapman & Hall, London, UK.

Saino, N., R. Ambrosini, M. Caprioli, A. Romano, M. Romano, D. Rubolini, C. Scandolara, and F. Liechti (2017). Sex-dependent carry-over effects on timing of reproduction and fecundity of a migratory bird. Journal of Animal Ecology 86: 239–249.

Seavy, N. E., D. L. Humple, R. L. Cormier, E. L. Porzig, and T. Gardali (2018). Evidence of the effects of climate change on landbirds in western North America: A review and recommendations for future research, in Trends and traditions: Avifaunal change in western North America (W. D. Shuford, R. E. Gill Jr., and C. M. Handel, Editors), pp. 331–343. Studies of Western Birds 3. Western Field Ornithologists, Camarillo, CA; doi 10.21199/SWB3.18.

Smith, M.A., N. Walker, I. Stenhouse, C. Free, M. Kirchhoff, O. Romanenko, S. Senner, N. Warnock, V. Mendenhall. 2014. A New Map of Important Bird Areas in Alaska. Anchorage: Audubon Alaska. https://sustainnorthdotorg.files.wordpress.com/2016/12/alaska_ibas_poster_withtext_6dec2014_mr.jpg

Smith, S. B., and P. W. Paton (2011). Long-term shifts in autumn migration by songbirds at a coastal eastern North American stopover site. The Wilson Journal of Ornithology 123: 557–566.

Stegman, L. S., Primack, R. B., Gallinat, A. S., Lloyd-Evans, T. L., and E. R. Ellwood (2017). Reduced sampling frequency can still detect changes in abundance and phenology of migratory landbirds. Biological Conservation 210: 107–115.

Stone, R. S., E. G. Dutton, J. M. Harris, and D. Longenecker (2002). Earlier spring snowmelt in northern Alaska as an indicator of climate change. Journal of Geophysical Research: Atmospheres 107(D10), ACL-10.

Sivakumar AH, Sheldon D, Winner K, Burt CS, Horton KG. 2021 A weather surveillance radar view of Alaskan avian migration. Proc. R. Soc. B 288: 20210232. 10.1098/rspb.2021.023

Tauzer, L.M. (2013). Recent changes in plant and avian communities at Creamer’s Refuge, Alaska using field and remote sensing observations. Master’s Thesis, University of Alaska Fairbanks, AK, USA. Tauzer_MSThesis_FINAL-1.pdf (usgs-cru-individual- data.s3.amazonaws.com)

Thompson, D. W., and J. M. Wallace (1998). The Arctic Oscillation signature in the wintertime geopotential height and temperature fields. Geophysical research letters 25: 1297–1300.

Tomotani B. M., P. Gienapp, D. G. M. Beersma, and M. E. Visser (2016). Climate change relaxes the time constraints for late-born offspring in a long-distance migrant. Proc. R. Soc. B 283: 20161366. 10.1098/rspb.2016.1366

Tonra, C. M., and M. W. Reudink (2018). Expanding the traditional definition of molt-migration. The Auk: Ornithological Advances 135: 1123–1132.

Visser, M. E. (2018). Climate change leads to differential shifts in the timing of annual cycle stages in a migratory bird. Global Change Biology 24: 823–835. 10.1111/gcb.14006

Tøttrup, A. P., K. Thorup, and C. Rahbek (2006). Changes in timing of autumn migration in North European songbird populations. Ardea 94: 527–536.

Trenberth, K. E., and J. W. Hurrell (1994). Decadal atmosphere-ocean variations in the Pacific.Climate Dynamics 9: 303–319.

Tulp, I., and H. Schekkerman (2008). Has prey availability for arctic birds advanced with climate change? Hindcasting the abundance of tundra arthropods using weather and seasonal variation. Arctic, pp.48–60.

US Fish and Wildlife Service (USFWS) (2015). Fall Migration Bird Banding: 20 Years of Monitoring Migratory Landbirds: summary, assessment and exploration of site move for Tetlin National Wildlife Refuge. U.S. Fish and Wildlife Service, Anchorage, AK. https://ecos.fws.gov/ServCat/Reference/Profile/74355

Urban, M.C., G. Bocedi, A. P. Hendry, J. B. Mihoub, G. Pe’er, A. Singer, J. R. Bridle, L. G. Crozier, L. De Meester, W. Godsoe, and A. Gonzalez (2016). Improving the forecast for biodiversity under climate change. Science, 353(6304), p.aad8466.

Van Buskirk, J., Mulvihill, R. S., & Leberman, R. C. (2010). Declining body sizes in North American birds associated with climate change. Oikos 119: 1047–1055.

Van Buskirk, J., R. S. Mulvihill, and R. C. Leberman (2009). Variable shifts in spring and autumn migration phenology in North American songbirds associated with climate change. Global Change Biology 15: 760–771.

Viereck, L. A., C. T. Dyrness, A. R. Batten, and K. J. Wenzlick (1992). The Alaska vegetation classification. Gen. Tech. Rep. PNW-GTR-286. Portland, OR: U.S. Department of Agriculture, Forest Service, Pacific Northwest Research Station. 278 p.

Vihma, T. (2014). Effects of Arctic sea ice decline on weather and climate: A review. Surveys in Geophysics 35: 1175–1214.

Visser, M. E., and C. Both (2005). Shifts in phenology due to global climate change: the need for a yardstick. Proceedings of the Royal Society B: Biological Sciences 272: 2561–2569.

Wallace, J. M., and D. S. Gutzler (1981). Teleconnections in the geopotential height field during the Northern Hemisphere winter. Monthly weather review 109: 784–812.

Wang, Y., H. Bi, H. Huang, Y. Liu, Y. Liu, X. Liang, M. Fu, and Z. Zhang (2019). Satellite- observed trends in the Arctic sea ice concentration for the period 1979–2016. Journal of Oceanology and Limnology 37: 18–37.

Ward, D. H., J. Helmericks, J. W. Hupp, L. McManus, M. Budde, D. C. Douglas, and K. D. Tape (2016). Multi-decadal trends in spring arrival of avian migrants to the central Arctic coast of Alaska: Effects of environmental and ecological factors. Journal of Avian Biology 47: 197–207.

Warnock, N. (2017). The Alaska WatchList 2017. Audubon Alaska, Anchorage, AK 99501.

Wells, J. V., editor. (2011). Boreal birds of North America: a hemispheric view of their conservation links and significance. University of California Press, Berkeley, California, USA.

Wells, J., D. Childs, F. Reid, K. Smith, M. Darveau, and V. Courtois (2014). Boreal Birds Need Half: Maintaining North America’s bird nursery and why it matters. Boreal Songbird Initiative, Seattle, Washington, Ducks Unlimited Inc., Memphis, Tennessee, and Ducks Unlimited Canada, Stonewall, Manitoba. http://www.borealbirds.org/sites/default/files/pubs/birdsneedhalf_0.pdf

Wendler, G., T. Gordon, and M. Stuefer (2017). On the precipitation and precipitation change in Alaska. Atmosphere, 8(12), p.253.

Wendler, G., L. Chen, and B. Moore (2012). The first decade of the new century: a cooling trend for most of Alaska. The Open Atmospheric Science Journal, 6(1).

Wilson, W. H. (2007). Spring arrival dates of migratory breeding birds in Maine: sensitivity to climate change. The Wilson Journal of Ornithology 119: 665–677.

Wu, J.X., B.L. Bateman, P.J. Hagelund, L. Taylor, A.J. Allstadt, D. Granfors, H. Westerkam, N.L. Mikel, C.B. Wilsey. 2022. U.S. National Wildlife Refuge System likely to see regional and seasonal species turnover in bird assemblages under a 2C warming scenario. Ornithological Applications 124: 1–14.

Zaifman, J., D. Shan, A. Ay, and A. G. Jimenez (2017). Shifts in bird migration timing in North American long-distance and short-distance migrants are associated with climate change. International Journal of Zoology. 10.1155/2017/6025646

Zelt, J., R. L. Deleon, A. Arab, K. Laurent, and J. W. Snodgrass (2017). Long-term trends in avian migration timing for the state of New York. The Wilson Journal of Ornithology 129: 271–282.

Zheng, J., Xu, X., & Jia, G. 2022. Effects of shifting spring phenology on growing season carbon uptake in high latitudes. Journal of Geophysical Research: Biogeosciences 127(12): e2022JG006900. 10.1029/2022JG006900

